# Mechanistic modeling of vascular tumor growth: an extension of Biot’s theory to hierarchical bi-compartment porous medium system

**DOI:** 10.1101/2020.06.29.176982

**Authors:** Giuseppe Sciumè

## Abstract

Existing continuum multiphase tumor growth models typically do not include microvasculature, or if present, this is modeled as non-deformable. Vasculature behavior and blood flow are usually non-coupled with the underlying tumor phenomenology from the mechanical viewpoint; hence, phenomena as vessel compression/occlusion modifying microcirculation and oxygen supply cannot be taken into account.

The tumor tissue is here modeled as a reactive bi-compartment porous medium: the extracellular matrix constitutes the solid scaffold; blood is in the vascular porosity whereas the extra-vascular porous compartment is saturated by two cell phases and interstitial fluid (mixture of water and nutrient species). The pressure difference between blood and the extra-vascular overall pressure is sustained by vessel walls and drives shrinkage or dilatation of the vascular porosity. Model closure is achieved thanks to a consistent non-conventional definition of the Biot’s effective stress tensor.

Angiogenesis is modeled by introducing a vascularization state variable, and accounting for tumor angiogenic factors and endothelial cells. Closure relationships and mass exchange terms related to vessel formation are detailed in a numerical example reproducing the principal features of angiogenesis. This example is preceded by a first pedagogical numerical study on one-dimensional bio-consolidation. Results are exquisite to realize that the bi-compartment poromechanical model is fully coupled (the external loads impact fluid flow in both porous compartments) and to envision further applications as for instance modeling of drugs delivery and tissue ulceration.

## 1 Introduction

Tumor growth is governed by a number of coupled phenomena, occurring at different spatial scales and often having very different characteristic times. It is a multiscale problem that involves various phases (cells, interstitial fluid, blood, extra-cellular matrix, etc.) and chemical species which interact biologically, chemically and also physically. Almost fifteen years ago, biologists began to address the limits of a purely molecular approach which produced a large amount of data that were not always clearly interpreted. Since then, researchers have gradually understood that physical forces in cancer microenvironment also have a first order effect on its proliferative-invasive behavior. Articles presenting this fresh perspective have been published in high impact journals, with very explicit titles such as: *“What does physics have to do with cancer?”* (1), *“Oncology: Getting physical”* (2), *“Mechanics: the forces of cancer”* (3), among others. Understanding that physics has a huge intrinsic and extrinsic impact on the evolution of cancer and its microenvironment, one strategy is to use physical sciences in the design and/or improvement of cancer therapies. With this aim, engineers, physicists and mathematicians are today paying growing attention to oncology, and mathematical models could play a pivotal role in deciphering the physics involved and its close relationship with cancer biology.

Proliferation and invasive behavior of tumor cells are influenced by anatomical substructures of afflicted organs (*e.g.* existing microvasculature, tissue-specific extracellular matrix, etc.) and their physical properties. The growing tumor compresses its surroundings, inducing a densification of the extracellular matrix (ECM) and stroma in the pathological zone. This reduces tissue permeability for drugs and nutrients ((1), (4)), while mechanical confinement pressure (exerted by the surrounding microenvironment) impacts proliferation rates and cell metabolism, and contributes to phenotype switch ((5), (3)). The tumor and induced strain significantly modify the physiological functions of the surrounding region because capillary vessels (ensuring blood microcirculation and oxygenation) are compressed as well as the lymphatic system. From a physical perspective, the disease can be viewed as a deregulation of the transport properties of the healthy tissue induced by coupled occurrence of several phenomena whose precise cause effect relationships are difficult to unravel. In fact, modification of the microenvironment induced by the tumor impacts the tumor itself since cells and their metabolism have to adapt to this evolving microenvironment (vessels compression, lack of oxygenation, stress, interstitial fluid accumulation, etc.). If we want to reproduce *in silico* such mechanisms to create a digital twin of cancer (useful to interpret associated phenomenology) biophysical modeling of micro-vascularization and surrounding co-opted vessels is required. This explains the very active research efforts aiming at modeling vascular tumor growth and angiogenesis (6; 7; 8). Different classes of modeling approaches exist in literature (continuous, mechanics-based models, cell-based models, and hybrid models, (9)) providing complementary descriptions of vascular network formation. Between these, macroscale multiphase models (sub-class of continuous models) emerge today as very promising approaches (10). Macroscale multiphase approaches enhance the consistency between the model and the real structure of the tumour microenvironment, enabling development of experimentally validated adjustments in contrast to modelling frameworks that treat the tumour as one continuous and homogeneous material.

Before going forward we shall start by clarifying the difference between a macroscale and a microscale description. Let us consider a virtual point, *P(x,y,z)*, within an tumor at the tissue level (hereafter also indicated as macroscale, see figure 1.a); then, upon magnification it with a microscope until small cell aggregates and single cells become visible, one discovers that the tissue architecture has several constituents, namely ECM, cells, blood vessels, etc. When we consider a point in the microscopy image a sole constituent is associated to this point, but on the other hand we have understood that behind the macroscopic point, *P(x,y,z)*, there is a Representative Elementary Volume (REV) of tissue where several constituents are present in a certain proportion, hereafter denoted as volume fraction (see figure 1). This overlapping of constituents is the basic concept of all existing macroscale multiphase models so also of the mathematical model presented in this paper. The great majority of existing multiphase model for tumor growth are founded on Mixture Theory (MT). Despite its large use, MT is exhaustive for modelling of particular *in vitro* configurations but not enough general to represent complexity and hierarchical structure of *in vivo* tumor tissue. The more recent Thermodynamically Constrained Averaging Theory (TCAT, (11)) is conversely able to physically and mathematically incarnate heterogeneity of tumor microenvironment. TCAT have two main advantages with respect to MT: *i)* by formally averaging the microscale equations up to the macroscale scale (instead of postulating them directly at the larger scale, as done in MT), larger scale variables are expressed precisely in terms of microscale precursors; *ii)* each phase may be constituted by several species. This second advantage allows us to make a rigorous distinction between: intra-phase transfer of mass/momentum (between species within the same phase) and inter-phase transfer of mass/momentum (from one phase to another one); advective and diffusive transport of species.

**Fig. 1.**
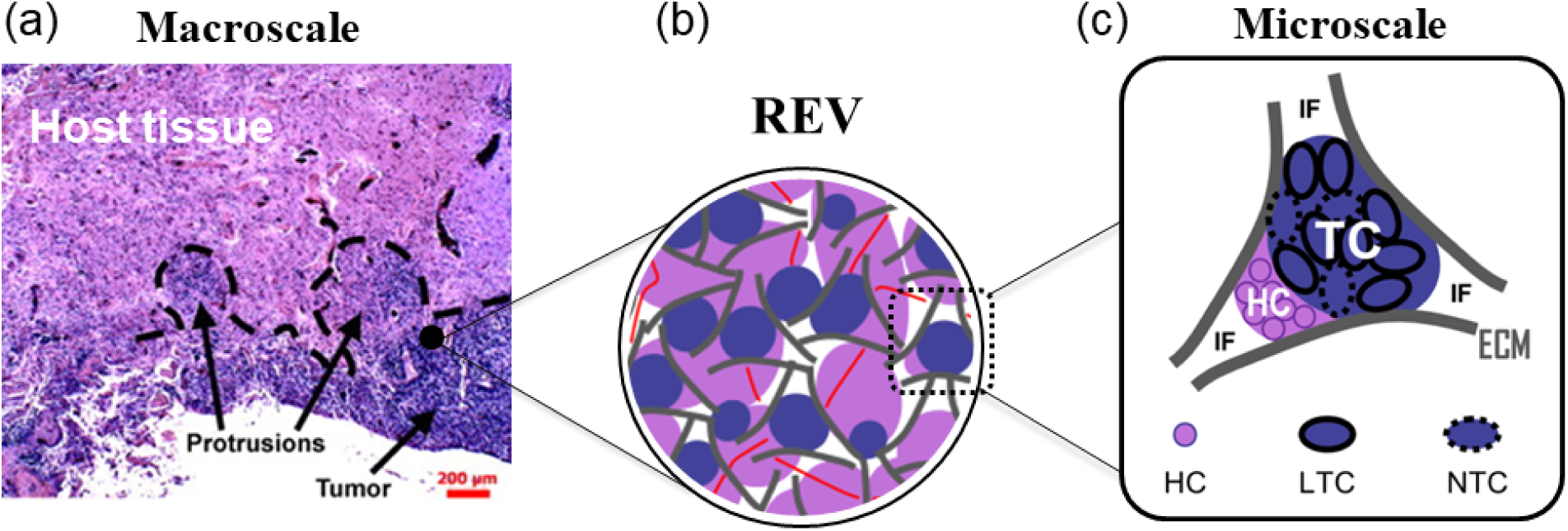
The multiphase system. (a) Infiltrative pattern from histopathology of human glioma (readapted from (16)); (b) representative elementary volume for the 5-phase system; (c) microscale configuration of phases.

TCAT provided us the theoretical background for the development of an original multiphase model for tumor growth during the avascular stage (12). This original version has been improved firstly considering a difference of pressure between tumor and healthy cells ((13), (14)), then removing the simplification assumption of a rigid ECM scaffold in (15). Starting from the existing multiphase model consisting of four phases (tumor cells, healthy cells, interstitial fluid and ECM), in this paper a major enhancement is presented to account for angiogenesis and the subsequent vascular stage of tumor growth; blood within micro-vasculature is modeled as an additional fluid phase. The vascular stage is preceded by a cascade of mechanisms. Tumor cells highly consume oxygen (due to their abnormal division rate and metabolism) so oxygen concentration within the cell aggregate decreases (especially in the internal areas which become quiescent or necrotic). Consequently, hypoxic cells start to produce tumor angiogenic factors (TAF) to stimulate formation of new blood capillaries and sprouting from existing vessels (angiogenesis) improving microcirculation and oxygen supply.

As it will emerge in the following paragraphs there are two big issues to develop a relevant biophysical model for angiogenesis and vascular growth.

The first one is the nonexistence in literature of some physiologically relevant (validated) constitutive relationships regulating dynamics of angiogenesis. Such constitutive equations are reasonably defined here (almost *ex novo*) for model closure. The proposed closure relationships have a general and qualitative significance; to gain a certain predictive potential these equations must be further specialized depending on the specific tumor of interest (cancer-specific model customization).

The second issue is coupling between ECM behavior and flow of four immiscible fluid phases (*i.e.* blood, interstitial fluid, tumor and host cell populations). The attention in the paper is mostly focused on this second issue because this is the most challenging from the modeling and computational viewpoint. A reliable mechanical coupling between the solid scaffold deformation, cell movement, interstitial fluid and blood flow is achieved thanks to an appropriate non-conventional definition of the Biot’s effective stress tensor and the ensuing derivation of a mechanical constitutive model accounting for vessel deformability and angiogenesis.

## 2 The Mathematical Model

Tumor tissue is modeled as a multiphase continuum so at the macroscopic level (tissue scale) at each point several phases coexist (each one characterized by its own volume fraction). Microscale conservation equations of mass and momentum are up-scaled *via* averaging theorems to obtain a larger scale (macroscale) set of equations. Within the averaging procedure all hypotheses/simplifications are explicitly introduced and macroscale variables are precisely defined. This ensures a clear connection between scales (11). The mathematical model allows for simulation of tumor growth in a vascularized tissue: the formulation presented in (15) for non-vascularized tumors is improved adding blood as a new fluid phase flowing within another porous compartment (representing micro-vasculature). This evolution is not trivial. A new definition of the effective stress tensor is needed to close the mathematical model and to couple the behavior of the five considered phases.

### 2.1 Tumor as a reactive multiphase continuum

The multiphase model comprises five phases: i) the tumor cells, *t*; ii) the healthy cells, *h*; iii) a solid scaffold, *s*; iv) the interstitial fluid, *l*; and v) blood, *b*. All other tissue constituents, as for instance chemical species (*e.g.* oxygen) and specific cell species (*e.g.* endothelial cells), are assumed to belong to one (or more) of the aforementioned phases.

The ECM, interstitial fluid (IF) and blood are present throughout the entire continuum domain, whereas tumor cells (TC) and healthy cells (HC) may exist or not, or exist only in certain subdomains. The solid scaffold consists essentially of extracellular matrix (ECM). All other phases are modeled as fluids. The TC phase consists of two dominant species: living tumor cells (LTC) and necrotic tumor cells (NTC). Necrosis is induced by low nutrient concentrations or excessive mechanical stress. The HC phase includes all non-pathologic cells of the multiphase system as for instance endothelial cells (EC) whose mass fraction is explicitly considerered in the model. The IF is a mixture of water and biomolecules, as nutrients, oxygen and waste products. Two species dissolved within the IF are explicitly considered: oxygen, *O*, which regulates tumor metabolism, proliferation and occurrence of necrosis; and a tumor angiogenic factor (TAF), *A*, produced by tumor cells and acting as a potent chemokine for endothelial cells. If we use the symbol, *ε*^*α*^, to indicate the volume fraction of the generic phase *α*, obviously the sum of the volume fractions for all phases gives the unit

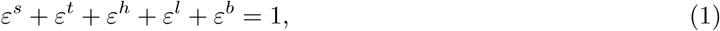

The system is modeled as a porous medium with two porous compartments. One part of pores, whose volume fraction is equal to *ε*^*t*^ + *ε*^*h*^ + *ε*^*l*^, is dominant and is saturated by cell populations and IF (phases *t, h* and *l*). The remains porous space is the capillary vessel porosity, *ε*^*b*^, filled by blood. In the following the porosity accessible to cells and IF is named “extra-vascular porosity” and indicated with *ε*

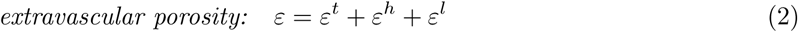

The porosity of the vascular compartment is named “vascular porosity”. This is saturated by blood so it corresponds to volume fraction of blood

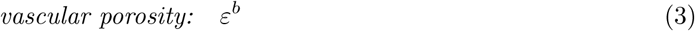

Blood walls have a solid nature but minor mass fraction and negligible “structural” functions at tissue level with respect to ECM scaffold.

Immiscibility between phases *t, h*, and *l*, is guaranteed by fluid-fluid interfacial tensions which can sustain differences in pressures between these three phases. Pressure differences between phases of the dominant extra-vascular porosity (*t, h* and *l*) and the blood in the vascular porosity is sustained by capillary vessels walls whose deformability is mechanistically considered and modeled.

### 2.2 Phases’ motion and material time derivatives

The description of phases’ motion is material for the solid phase and spatial for the fluid phases whose motion is refereed to that of the solid phase. In other words, the deforming solid scaffold is the reference space where fluid phases’ motion is described in an Eulerian way. This approach is customary in geomechanics (17). Independently on the employed description, conservation equations must refer to the current position of the multiphase continuum and hence must primarily be formulated using a spatial description as presented in the next paragraph. The equations are then expressed in a material form introducing material time derivatives with respect of the deforming solid scaffold. The material time derivative of any differentiable function, *f*^*π*^, given in its spatial description and referring to a moving particle of the *π* phase, is

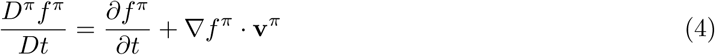

More in general, we can also express a material derivative with respect of the movement of another phase *α* (the time derivative is taken moving with the phase *α*)

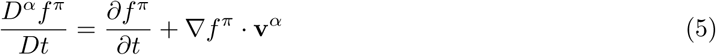

Subtracting the previous two eqs yields the following relation

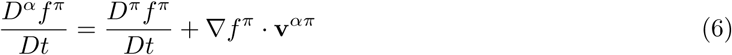

where

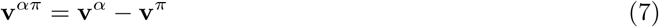

is commonly called relative velocity, and is the velocity of the phase *α* with respect to the phase *π*.

### 2.3 General form of governing equations from TCAT

In this section conservation equations for mass and momentum (in their general spatial form) are reported and explained (11). These equations are subsequently specialized for the analyzed multiphase system in the next subsection.

The spatial form of the mass balance equation for an arbitrary phase *α* reads

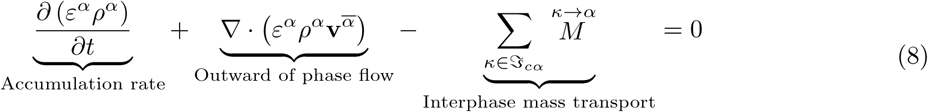

where *ρ*^*α*^ is the density, 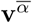 is the local velocity vector, 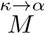 are the mass exchange terms accounting for transport of mass at the *κα* interface from phase *κ* to phase *α*, and 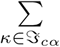 is the summation over all the phases sharing interfaces with the phase *α*.

An arbitrary species *i* dispersed within the phase *α* has to satisfy mass conservation too. The following spatial equation is derived following TCAT

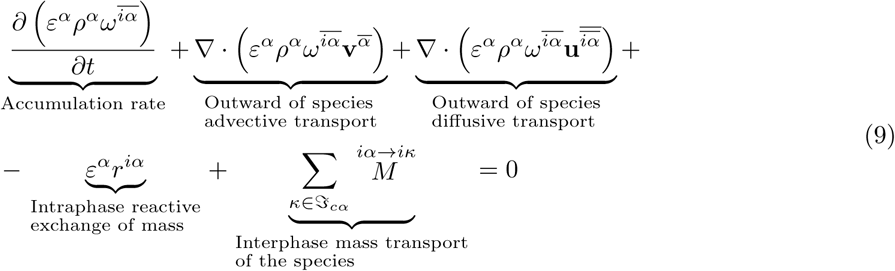

where 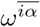 identifies the mass fraction of the species *i* dispersed with the phase *α, ε*^*α*^*r*^*iα*^ is a reaction term that allows to take into account the reactions between the species *i* and the other chemical species dispersed in the phase *α*, and 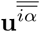 is the diffusive velocity of the species 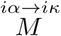 are mass exchange terms accounting for mass transport of species *i* at the *κα* interface from phase *α* to phase *κ*. Applying the product rule and introducing eqn (8), the previous equation can be also written in this alternative form (commonly called distribution form)

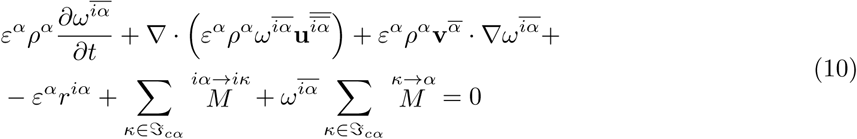

Concerning momentum conservation, given the characteristic time scale of the modeled problem and the assumption of quasi-static processes, inertial forces can be neglected as well as forces due to mass exchange due to the small difference in density between cells and aqueous solutions. This allows simplifying the form of the linear momentum balance equation provided by TCAT for a general phase *α* which, neglecting also gravitational body forces, becomes (12)

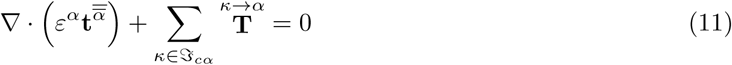

where 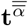 is the stress tensor of the phase *α* and 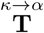 is the interaction force between phase *α* and the adjacent phases. Eqn (11) which applies for a generic phase *α* (solid or fluid) can be expressed in an alternative form for fluid phases. Indeed, for relatively slow flow, the stress tensor for a fluid phase, *f*, can be properly approximated as

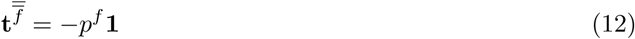

where *p*^*f*^ is the averaged fluid pressure and **1** the unit tensor. Then, from TCAT – see appendix A, of (12) – it can be shown that the momentum balance equation for fluid phases can also be expressed as

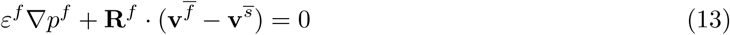

where **R**^*f*^, is a symmetric second order resistance tensor accounting for interaction between the fluid phase, *f*, and the solid phase, *s*.

### 2.4 Governing equations of the multiphase tumor tissue model

The model is governed by conservation equations of mass and momentum of considered phases and species. Such equations are detailed in the following dedicated paragraphs.

#### 2.4.1 Mass conservation equations (11 scalar independent eqs)

From the definition of extra-vascular porosity, *ε*, (eqn (2)), and considering the constraint eqn (1) it follows that

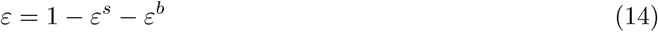

Considering now the extra-vascular fluid phases (*t, h* and *l*) and defining their own saturation degree as *S*^*β*^ = *ε*^*β*^ *ε* (with *β* = *t, h, l* index associated to extra-vascular fluids), another constraint equation (alternative to eqn (1)) can be rewritten in term of saturation degrees

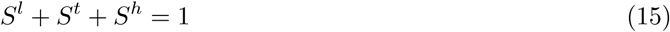

It is emphasized that such saturation degrees refer to the extra-vascular porosity, *ε*, only. Using eqn (5) to express derivatives with respect of the moving solid phase and introducing extra-vascular porosity and saturation degrees of *t, h*, and *l*, mass balance equations of *s, t, h, l* and *b* phases read respectively

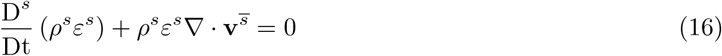

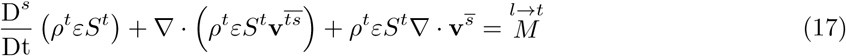

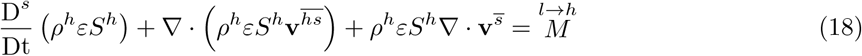

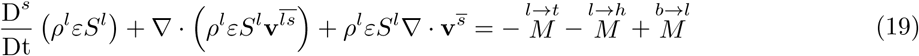

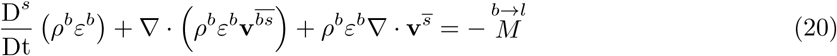

where 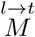 is the mass exchange from IF to the tumor due to cells growth and metabolism, 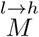 is the mass exchange from IF to the HC population, and 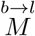 allows to account for exchange of mass between blood and IF.

The tumor cell phase is modeled as a mixture of living and necrotic cells containing also water, *W*, oxygen, *O*, TAF, *A*, and other chemical species in minor proportion (18). Evolution of mass fraction of necrotic tumor cell species, 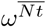, is here modeled by considering its mass conservation equation. From eqn (9), assuming that there is no diffusion of necrotic cells and introducing the material derivative with respect of the solid phase, mass conservation equation of necrotic cell species becomes

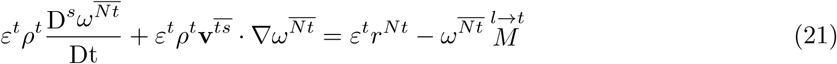

where *ε*^*t*^*r*^*Nt*^ is the rate of necrosis of tumor cells.

HC phase consists of all non-pathological cells of tumor microenvironment and, similarly to the tumor cell phase, also includes water, *W*, oxygen, *O*, TAF, *A*, and other chemical species in minor proportion. Hence, EC species (endothelial cells are elementary constituents to form the microvascular system) belongs to HC, phase *h*, and move within it due to advection-diffusion. The EC conservation equation reads

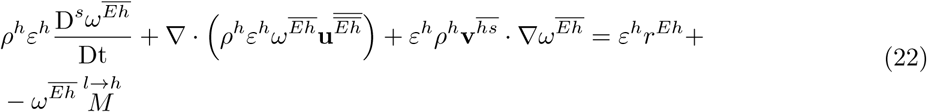

where *ε*^*h*^*r*^*Eh*^ is an intra-phase exchange of mass accounting for EC production due to reaction of water with other chemical species (contained in the phase *h*). Here it is assumed that mass provided by this reaction corresponds to the overall mass exchanged with IF, 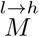, which is instantaneously used to produce 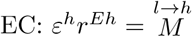. This assumption allows us to rewrite the previous equation as

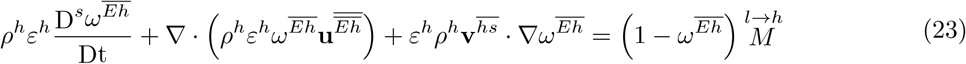

IF is a mixture of water (solvent) and several diluted chemical species. Oxygen, *O*, and TAF, *A*, are between these dilute IF species: evolution of their relative mass fraction is governed by two additional mass conservation equations

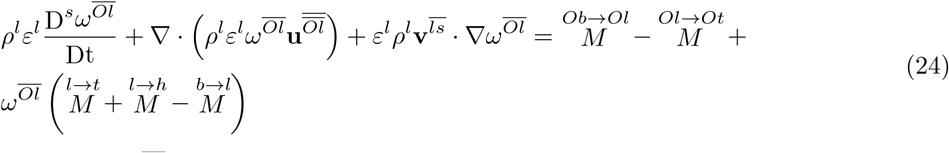

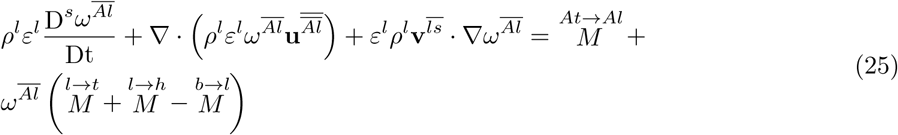

where 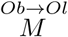 is oxygen provided by blood to IF, 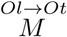 is oxygen consumed by tumor cells do to their metabolism and proliferation rate, 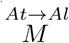 are TAF released by tumor cells.

#### 2.4.2 Linear momentum conservation equations (15 scalar independent eqs)

Summing eqn (11) over all phases gives the momentum equation of the whole multiphase system as

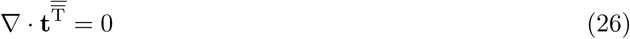

where 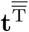 is the total Cauchy stress tensor acting on the multiphase system. For convenience eqn (26) is used instead of the linear momentum balance equation of the solid phase, so the variable 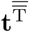 is introduced at the place of 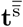 (if needed this last can be computed from other primary variables of the model).

The alternative equivalent form (*i.e.* eqn (13)) is considered for fluid phases. In porous media mechanics typically eqn (13) is expressed as

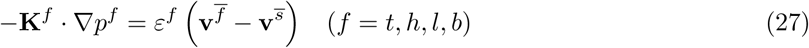

where **K**^*f*^ is called hydraulic conductivity. The hydraulic conductivity depends on the properties of both the flowing fluid and the solid porous material *via* the fluid dynamic viscosity and the intrinsic permeability of the solid matrix respectively. For blood, momentum eqn (27) is valid under the hypotheses of slow laminar flow with negligible inertial effects. A more general relationship including inertial effect and impact of the deformation rate, 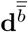, may be preferable under certain situations.

#### 2.4.3 Model closure strategy: summary of model variables versus conservation equations

In the previous two paragraphs 26 scalar independent equations have been presented for a total number of 49 scalar independent variables; this means that 23 scalar constitutive relationships are needed to close the mathematical model. To reduce the complexity of the mathematical model the following simplification hypothesis is introduced:

##### Hypothesis 1

*Compressibility of phases is assumed having a minor impact on system evolution. Hence, phases’ densities are assumed constant and phases Bulk’s moduli very large with respect to the average Bulk’ modulus, K, of the porous scaffold:* 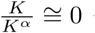 *with α* = *s, t, h, l, b.*

Assuming constant phases’ density allows reducing the number of independent variables to 44 (see table 1), so 18 constitutive relationships are needed namely: a state equation for volume fraction of the blood phase, (1 scalar eqn); the solid scaffold mechanical constitutive model (6 scalar eqs); pressures-saturations relationships for pressure difference between fluids *t, h*, and *l* in the extra-vascular porosity (2 scalar eqs, (13)); Fick type laws for diffusive velocity of oxygen, TAF and EC species (9 scalar eqs).

**Table 1.** Summary of the model independent variables.

| Phases | Ind. | Species | Modeled species | Associates variables | Scalar variables |
| --- | --- | --- | --- | --- | --- |
| ECM | $s$ | - | - | $\varepsilon^s, \varepsilon, \mathbf{t}^{\overline{\overline{T}}}, \mathbf{v}^{\overline{\overline{s}}}$ | 11 |
| Tumor cells | $t$ | $L, N, W, O, A$ | $Nt$ | $S^t, p^t, \mathbf{v}^{\overline{\overline{t}}}, \omega^{\overline{\overline{Nt}}}$ | 6 |
| Host cells | $h$ | $E, W, O, A$ | $Eh$ | $S^h, p^h, \mathbf{v}^{\overline{\overline{h}}}, \omega^{\overline{\overline{Eh}}}, \mathbf{u}^{\overline{\overline{Eh}}}$ | 9 |
| Int. fluid | $l$ | $W, O, A$ | $Oh, Ah$ | $S^l, p^l, \mathbf{v}^{\overline{\overline{l}}}, \omega^{\overline{\overline{Ol}}}, \omega^{\overline{\overline{TAFI}}}, \mathbf{u}^{\overline{\overline{Ol}}}, \mathbf{u}^{\overline{\overline{TAFI}}}$ | 13 |
| Blood | $b$ | $W, O$ | - | $\varepsilon^b, p^b, \mathbf{v}^{\overline{\overline{b}}}$ | 5 |
| Tot. number of scalar unknowns |  |  |  |  | 44 |

### 2.5 Angiogenesis, effective stress tensor and constitutive relationships

In this paragraph the angiogenesis model is introduced together with definition of effective stress tensor and other constitutive relationships needed for model closure.

#### 2.5.1 Angiogenesis rate function

Angiogenesis is modeled thanks to the introduction of an internal variable, *Γ*, describing advancement of vessel formation. *Γ* = 0 corresponds to a normal microvasculature while *Γ* = 1 corresponds to a fully developed abnormal vasculature (when angiogenesis is ended). Multiple could be the assumptions about the functional dependencies of the internal variable, *Γ*. Very frequently in literature it is assumed that the density of capillary vessels is proportional to the density of endothelial cells (see for instance (19; 7)). Here a similar assumption is adopted: angiogenesis is initially proportional to the increase of EC mass fraction; however, with advancement of vessel formation, this proportionality progressively decreases thanks to a factor (1 *− Γ*). The following rate relationship is proposed

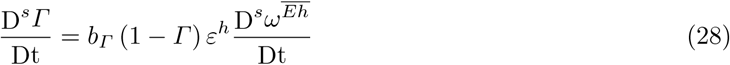

where *b*_*Γ*_ is constant parameter governing the rate of vessel generation.

#### 2.5.2 Effective stress tensor: a non-conventional form

The solid phase stress tensor, 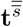, can be decomposed into component parts (11)

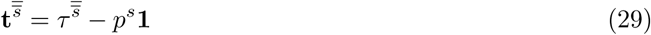

where 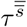 is an effective solid phase stress tensor and *p*^*s*^ is the solid phase pressure. From TCAT the total stress tensor, 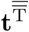, is the summation of all phase tensors each one weighed *via* its own volume fraction

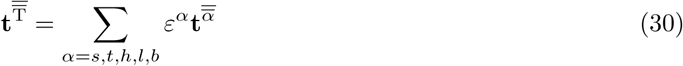

Introducing eqs (29) and (12) in eqn (30) gives

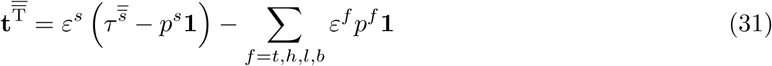

The following two hypotheses are introduced:

##### Hypothesis 2

*Blood vessels are mostly surrounded by extra-vascular fluids, TC, HC and IF, so they have weak direct mechanical interaction with the solid ECM scaffold.*

##### Hypothesis 3

*Consistently with hypothesis 2 the solid pressure, p*^*s*^, *is assumed being related to pressures of extra-vascular fluids only (phases t, h and l):*

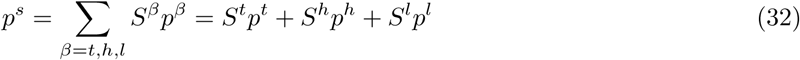

Introducing eqn (32) in eqn (31), and using eqn (14) give the total stress tensor as

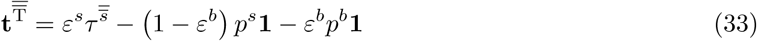

so, rearranging the terms, the effective stress tensor, 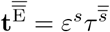, under introduced hypotheses, reads

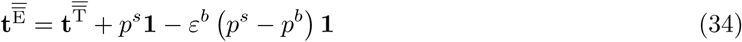

#### 2.5.3 State equation for volume fraction of blood

Consistently with hypothesis 2, it can be reasonably assumed that volume fraction of blood vessels, *ε*^*b*^, depends on the difference between the blood pressure inside the vessels *p*^*b*^ and an average external pressure exerted by extra-vascular fluid phases. Such extra-vascular pressure is assumed equal to the solid pressure, *p*^*s*^ (eqn (32)). The following relationship is proposed for *ε*^*b*^

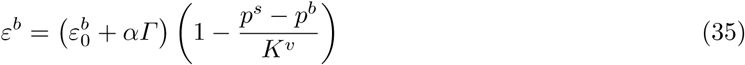

where 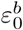 is the volume fraction when *p*^*s*^ = *p*^*b*^ and without angiogenesis (*i.e.* when *Γ* = 0), *K*^*v*^ is the vessel compressibility and *α* is a constant parameters representing the additional vasculature for a fully developed angiogenesis (*i.e.* when *Γ* = 1).

#### 2.5.4 Mechanical constitutive model

The overall Bulk’s modulus of the solid scaffold is negligible with respect of that of the solid phase (hypothesis 1) so the Biot’s coefficient equals 1 and the effective stress tensor, 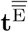 of eqn (34), is the stress responsible for all deformation of the solid ECM scaffold. Assuming a linear elastic behavior we can write a first form of the mechanical constitutive model as

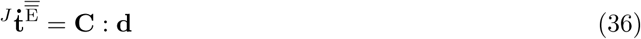

where 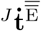 is the Jaumann objective time derivative of the effective stress tensor, **C** is the elasticity tensor and **d** is the strain rate. From eqn (34) Jaumann stress rate of effective stress must be also given by

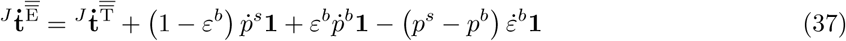

The mechanical constitutive relationship can therefore be expressed also in term of the Jaumann rate of the total stress tensor. Simple operations give

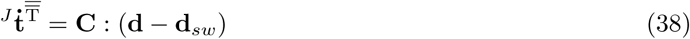

with **d**_*sw*_ isotropic swelling strain rate given by

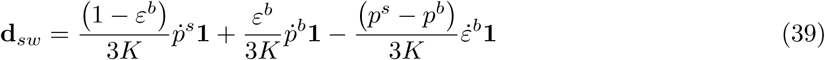

*K* is the Bulk’s modulus of the solid porous scaffold (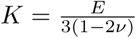, *E* and *ν* being Young’s modulus and Poisson’s ratio respectively). If 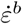 is expressed from variation with time of variables *p*^*s*^, *p*^*b*^ and *Γ*, some calculations allow rewriting the isotropic swelling strain rate **d**_*sw*_ as

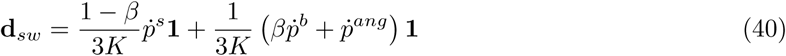

with *β* compressibility function and 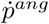 angiogenesis pressure rate equal respectively to

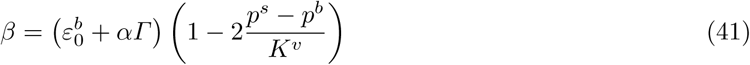

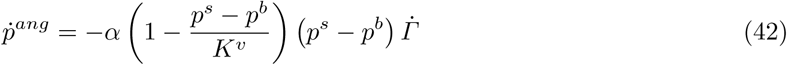

A hierarchy of models can be defined from eqs (38) and (40) (see Figure 2). The most general constitutive equation is for vascular growth with angiogenesis (VGA)

**Fig. 2.**
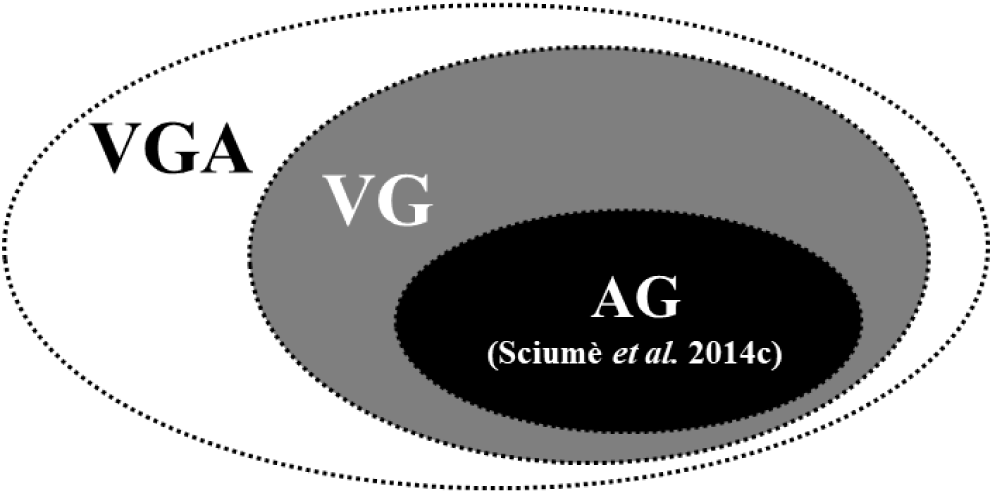
Model hierarchy: vascular growth model with angiogenesis (VGA); vascular growth model (VG); avascular growth model (AG).

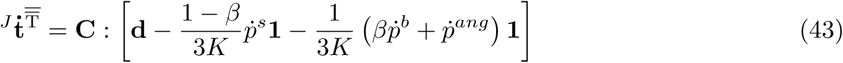

If angiogenesis is not considered, 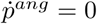, so the previous equation simplifies for vascular growth (VG) as

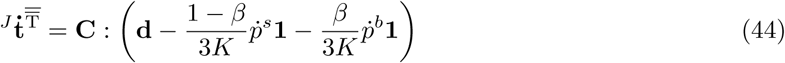

with

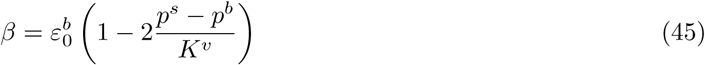

Finally if neither angiogenesis and vasculature are considered, 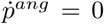 and *β* = 0, so the equation reduces to that used in (15) for the avascular growth model (AG)

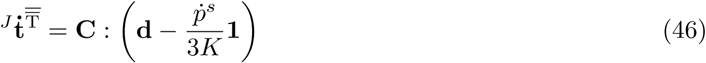

#### 2.5.5 The two pressures-saturations relationships

Being the extravascular porosity saturated by three immiscible fluid phases, and having each phase its own pressure three capillary pressures *p*^*ij*^ (pressure difference between *fluid*_*i*_ and *fluid*_*j*_) can be defined

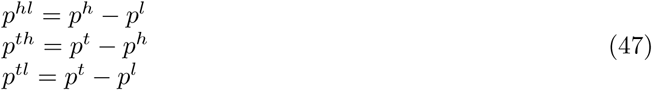

As in (13) and in (14) we assume here that IF is the wetting fluid, HC is the intermediate-wetting fluid and TC the non-wetting one. Only two between the previously defined capillary pressures are independent since

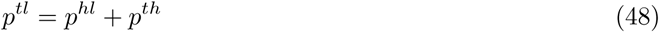

The two capillary pressure-saturation relationships read (13)

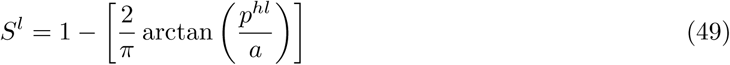

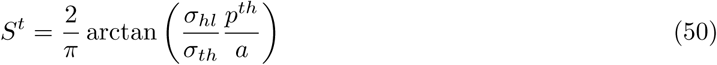

where *a* is a constant parameters depending on ECM microstructure, *σ*_*hl*_ and *σ*_*th*_ are HC-IF and TC-HC interfacial tension respectively. Using definition of capillary pressures, the solid pressure, eqn (32), can also be expressed as

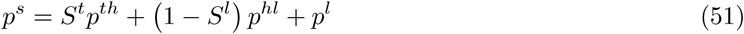

#### 2.5.6 Diffusive velocities of oxygen, TAF and EC

To approximate the diffusive flux in eqs (24) and (25), Fick’s law is used

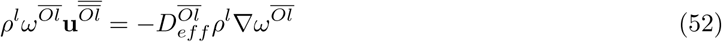

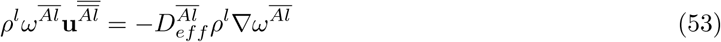

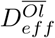 and 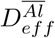 are the effective diffusion coefficient of oxygen and TAF respectively. The effective diffusion coefficient is a function of IF volume fraction

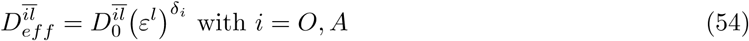

where 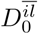 is the diffusion coefficient of species *i* in the bulk interstitial fluid and *δ*_*i*_ is a constant coefficient. Actually, effective diffusion coefficients have a nonlinear dependence on the volume fraction of IF, because diffusivity depends on the connectivity grade of the extra-cellular spaces (*interstitium*). *δ*_*i*_ is set equal to 2.

To account for diffusion of endothelial cell species a more complex closure relationship is adopted. This contains a fickian diffusion term and a coupling term to account for *chemotaxis* induced by TAF gradient. Coupling with TAF gradient is achieved thanks to a constant coefficient *C* identified numerically

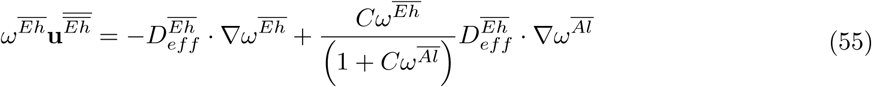

Details about derivation of eqn (55) are reported in the Appendix A.

## 3 Numerical applications

Two numerical applications are presented in the following paragraphs.

The first one is a one-dimensional consolidation case of a bi-compartment porous medium in which each compartment is saturated by a sole fluid: blood for the vascular porosity and a relatively viscous interstitial fluid (with dynamic viscosity *µ*^*l*^ = 1 Pa.s) for the extravascular porosity. This academic case, thanks to its low degree of complexity (the presented mathematical model reduces importantly, because of the non-reactive nature of the system and the absence of cell populations), allows a clear interpretation of system behavior.

In the second example the full model is applied to simulate an initial avascular tumor growth within two vascular beds followed by stimulation of angiogenesis and increase of the degree of vascularization of the tissue.

For both numerical examples once the application presented (geometry, initial and boundary condition), the final form of the governing equations is derived, then numerical results are presented.

### 3.1 One-dimensional bio-consolidation: one fluid + blood

The first analyzed case is the simplest configuration of bi-compartment porous medium: one fluid in the extra-vascular porosity and blood within capillary vessels. This example is essential to understand how the model works and how flow of blood and the extravascular fluid are coupled. A column of 0.1 mm of height is compressed on the top with a load going from 0 Pa to *p*_0_ = 200 Pa in 5 seconds, then sustained at 200 Pa during 120 seconds. The applied load increases following a sinusoidal ramp (see figure 3 on the right). This facilitates convergence of the time integration scheme. Time labelled as zero correspond to the time when the full load, *p*_0_, is applied (so the numerical simulation start 5 seconds before). This choice has been done to compare the obtained numerical solution with the analytical one of the simple porosity case (*i.e.* the Terzaghi problem) available for a load applied instantaneously. Such analytical solution gives the interstitial fluid pressure as series expansion function of the 1D-coordinate, *z* and time, *t*.

**Fig. 3.**
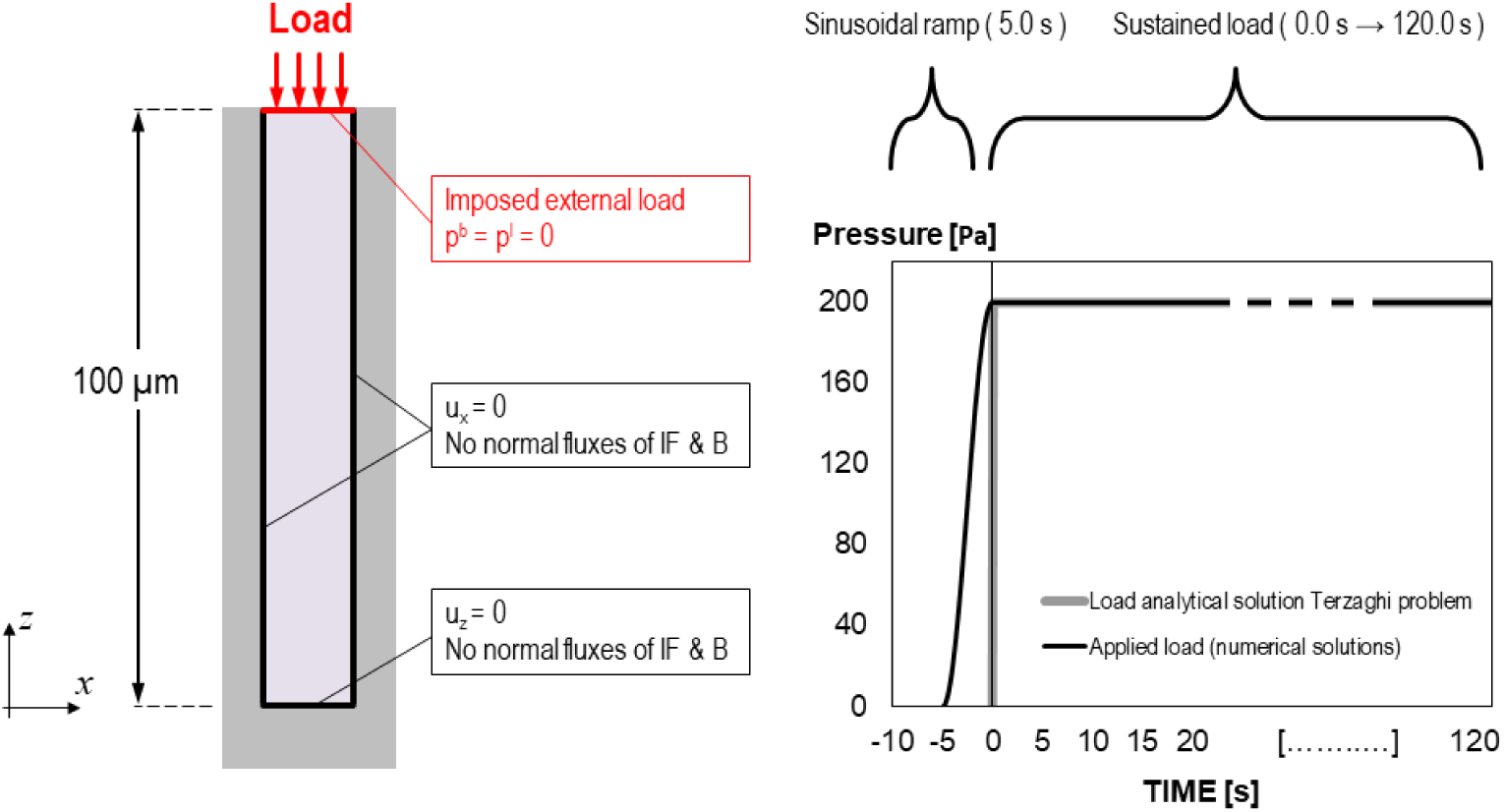
Geometry and boundary conditions of the modelled case (left); sinusoidal rump modeling the load: time “0” correspond to the time when the full load is applied (right).

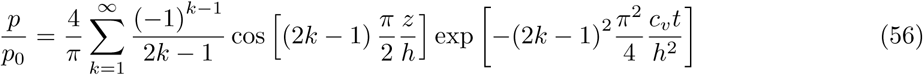

Where *c*_*v*_ is the consolidation coefficient

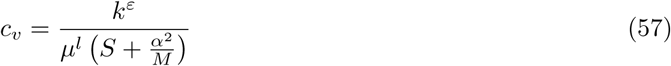

In eqn (57) *k*^*ε*^ is intrinsic permeability of the extra-vascular porosity, *µ*^*l*^ is dynamic viscosity of the extravascular fluid, *M* is the longitudinal modulus (or also called constrained modulus), 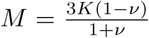,*α* is the Biot’s coefficient, and *S* is the inverse of Biot’s modulus

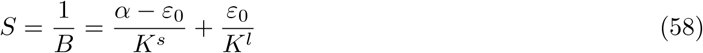

As assumed (hypothesis 1) phases’ compressibility is very larger with respect to the overall porous medium compressibility. This allows assuming Biot’s coefficient, *α*, equal to 1 and S = 0. Consequently the consolidation coefficient simplify to

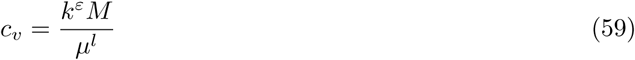

Concerning boundary conditions at the top edge (red line in the left part of figure 3) Dirichlet’s conditions are imposed for pressures of fluid phase (*p*^*l*^ = *p*^*b*^ = 0 Pa). On the rest of the boundary (black lines in the left part of figure 3) no fluid flow is permitted (*i.e.* sealed condition), horizontal displacement is restrained for the vertical bounds and vertical displacement is restrained for the bottom bound.

#### 3.1.1 Derivation of the final form of governing equations

The primary variables of the model are *p*^*l*^, *p*^*b*^ and **u**^*s*^. Interstitial fluid and blood are considered as a pure phases. Furthermore no mass exchange terms are present. The presented system of governing equations reduce substantially as follows (note that *ε*^*l*^ = *ε* and therefore *S*^*l*^ = 1)

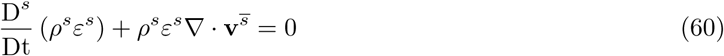

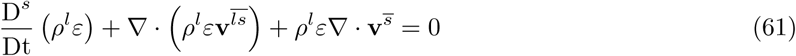

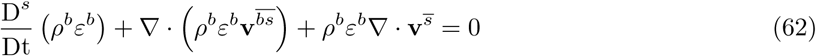

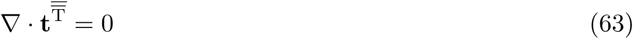

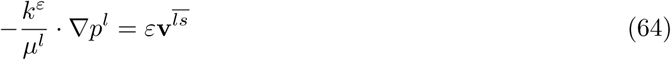

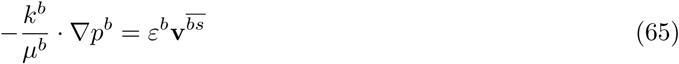

where 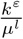 and 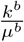 are hydraulic conductivity of interstitial fluid and blood phases respectively. Dividing eqn (60) for *ρ*^*s*^ and introducing eqn (14), some simple operations give

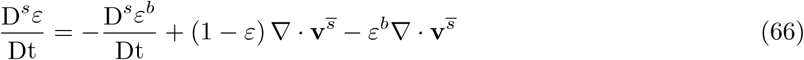

The mass balance equation of interstitial fluid is now considered. Dividing it for *ρ*^*l*^, and exploiting the product rule, eqn (61) can be rewritten as

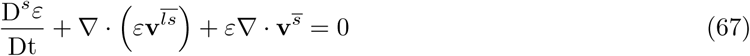

Introducing eqn (66) and (64) in eqn (67) gives

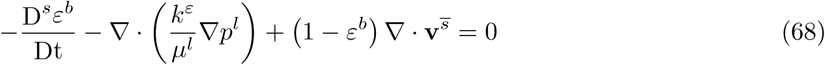

Performing analogous operations eqn (62) can be rewritten as

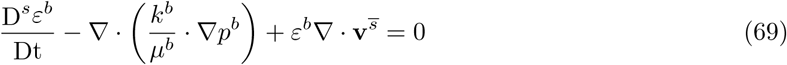

Thinking forward on numerical solution, it is useful to express eqs (68) and (69) in term of temporal variation of primary variables *p*^*b*^ and *p*^*l*^. From state equation (35), disregarding angiogenesis term, and being in this case *p*^*s*^ = *p*^*l*^ the time variation of volume fraction of blood vessels can be computed as

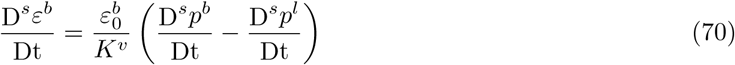

Introducing eqn (70) in eqs (68) and (69) and expressing eqn (63) in rate form (customary in geomechanics for incremental numerical resolution, see (17)) the governing final system of coupled equations reads

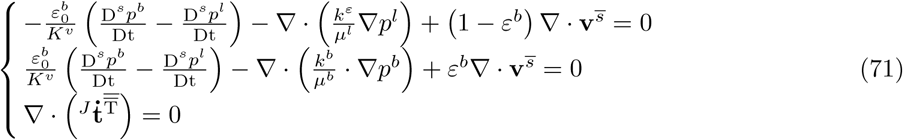

All parameters used for the numerical example are reported in table 2. The weak form of governing equations is obtained by means of the standard Galerkin procedure and is then discretized in space by means of the finite element method. Integration in the time domain is carried out with the generalized mid-point rule where a Crank-Nicolson scheme is used. The fix point method is used to account for the nonlinear nature of the system of equations. A mixed finite element space is adopted where quadratic elements (triangular with six nodes and quadrangular with eight nodes) are used for **u**^*s*^, while linear elements (triangular with three nodes and quadrangular with four nodes) are used for *p*^*l*^ and *p*^*b*^. The described model has been implemented in the code CAST3M (http://www-cast3m.cea.fr) of the French Atomic Energy Commission.

**Table 2.** Summary of input parameters for the three modelled cases.

| Parameter class | Parameter | Symbol | Value | Unit |
| --- | --- | --- | --- | --- |
| ECM and<br>extravascular<br>porosity | Young's modulus | $E$ | 5000 | $Pa$ |
| | Poisson's ratio | $\nu$ | 0.2 | - |
| | Porosity | $\varepsilon$ | 0.5 | - |
| | Intr. permeability | $k^\varepsilon$ | 1.E-14 | $m^2$ |
| | Dynamic viscosity IF | $\mu^l$ | 1.0 | $Pa.s$ |
| Vascular<br>porosity<br>*Case1* | Initial vessel fraction | $\varepsilon_0^b$ | 0.02 | - |
| | Vascular permeability | $k^b$ | 2.E-16 | $m^2$ |
| | Vessels Bulk's modulus | $K^v$ | 1000 | $Pa$ |
| | Blood dynamic viscosity | $\mu^b$ | 0.004 | $Pa.s$ |
| Vascular<br>porosity<br>*Case2* | Initial vessel fraction | $\varepsilon_0^b$ | 0.04 | - |
| | Vascular permeability | $k^b$ | 4.E-16 | $m^2$ |
| | Vessels Bulk's modulus | $K^v$ | 1000 | $Pa$ |
| | Blood dynamic viscosity | $\mu^b$ | 0.004 | $Pa.s$ |

#### 3.1.2 Results

Figure 4 shows the extravascular pressure profile along the one dimensional coordinate at three different times: 20 seconds, 50 seconds and 80 seconds after the application of the full load. Three cases are studied:

**Fig. 4.**
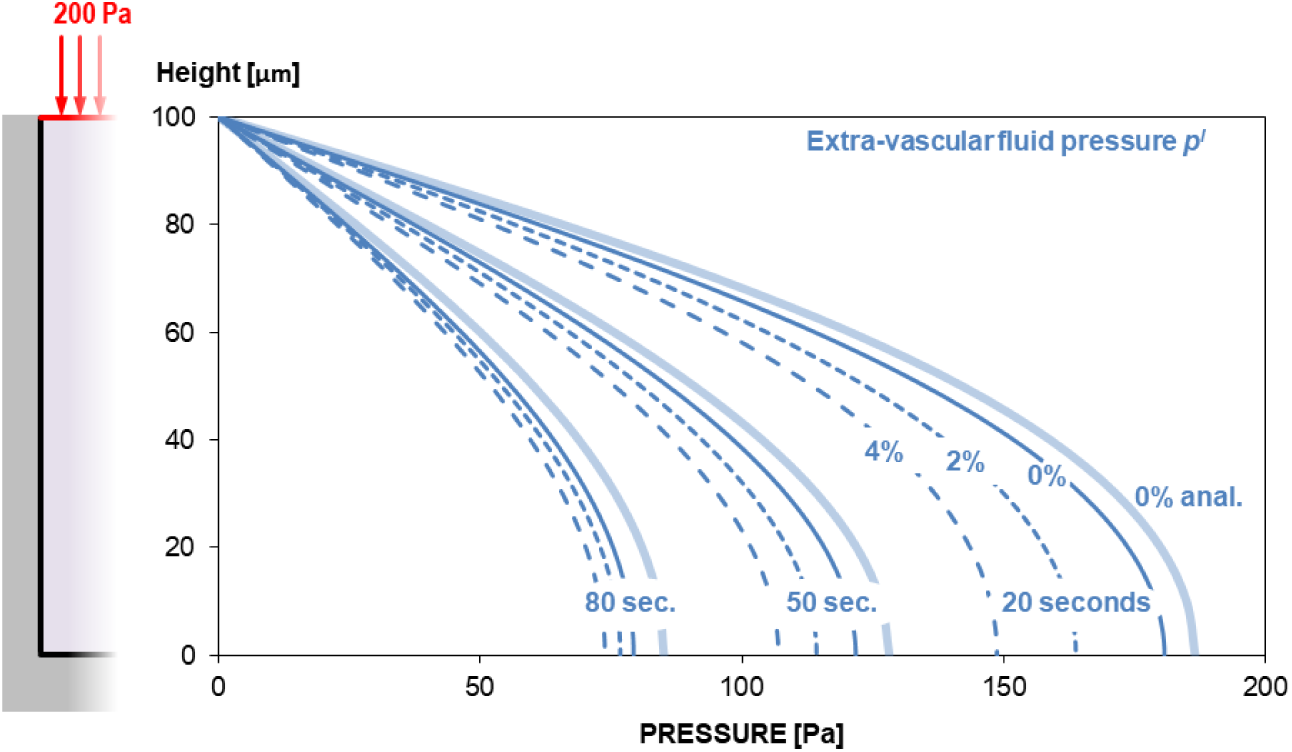
Extra-vascular fluid pressure, *p*^*l*^ along the one-dimensional coordinate for the three modeled cases.

– “0%” is the case with no vascular porosity, this corresponds with a porous medium saturated by the sole interstitial fluid; for this case the analytical solution calculated with eqn (56) is provided;
– “2%” is the case with a volume fraction of blood initially equal to 0.02;
– “4%” is the case with a volume fraction of blood initially equal to 0.04.

A difference can be appreciated between the numerical and analytical solution for the 0% case. This is due to difference in the application of the load that in the numerical case is sinusoidal and starts 5 second before the instantaneous load assumed for the analytical solution (see figure 3, right). As can be appreciated by the graphs the higher the volume of vessel is the lower will be the amplitude of the extravascular fluid pressure. Actually, vessels deformation contribute to dissipate more rapidly the increase of extravascular pressure induced by the application of the load. Figure 5.a shows the displacement of the top point (z = 100 *µm*) whose initial amplitude increases with the increase of the vascular porosity. Despite the initial differences the vertical displacement after a certain time tends asymptotically to the same value for the three analyzed cases. Figure 5.b shows evolution of blood and extravascular fluid pressures versus time at the bottom point (z = 0 *µm*). This graph another time shows the effect that vasculature has on reducing the extravascular fluid pressure. From figure 5.b one can also observe that the difference on initial blood volume fraction does not affect importantly the peak value of blood pressure. It is interesting to note that the blood pressure becomes negative 5 second after the full application of the load. In the first phase (from −5 to 5 seconds) blood is expulsed because part of the deformation is absorbed by the vasculature which reduces its volume; then the extravascular pressure decreases slowly so the pressure on vessel walls also decreases and vascular porosity regains gradually its initial volume fraction (blood comes back inside the vasculature). This behavior can be also appreciated in figures 5.c and 5.d where evolution of extravascular and vascular porosity at the bottom point (z = 0 *µm*) are depicted. A counterpart of the volumetric deformation is initially taken by vessels, then, with the decrease of interstitial fluid pressure, the volume fraction of vessels regains progressively its initial value whereas extravascular porosity decreases with time due to consolidation. In figure 6 it is possible to understand schematically how the load is transferred from the solid scaffold to the vascular system by mediation of the extravascular fluid.

**Fig. 5.**
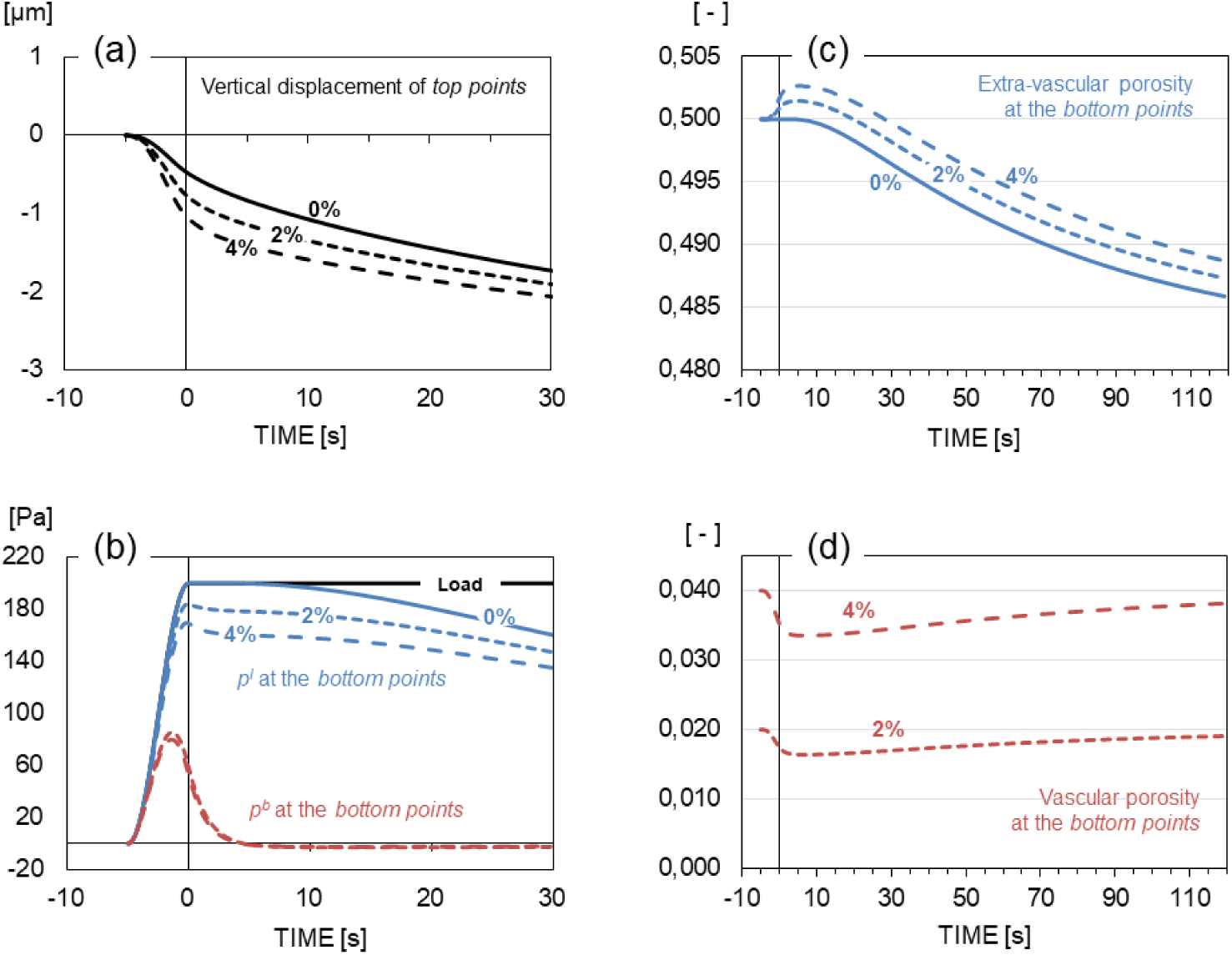
Evolution of the solution with time at fixed points: (a) vertical displacement at the top of the column; (b) extravascular fluid pressure and blood pressure at the bottom of the column; (c) and (d) extravascular and vascular porosities at the bottom of the column.

**Fig. 6.**
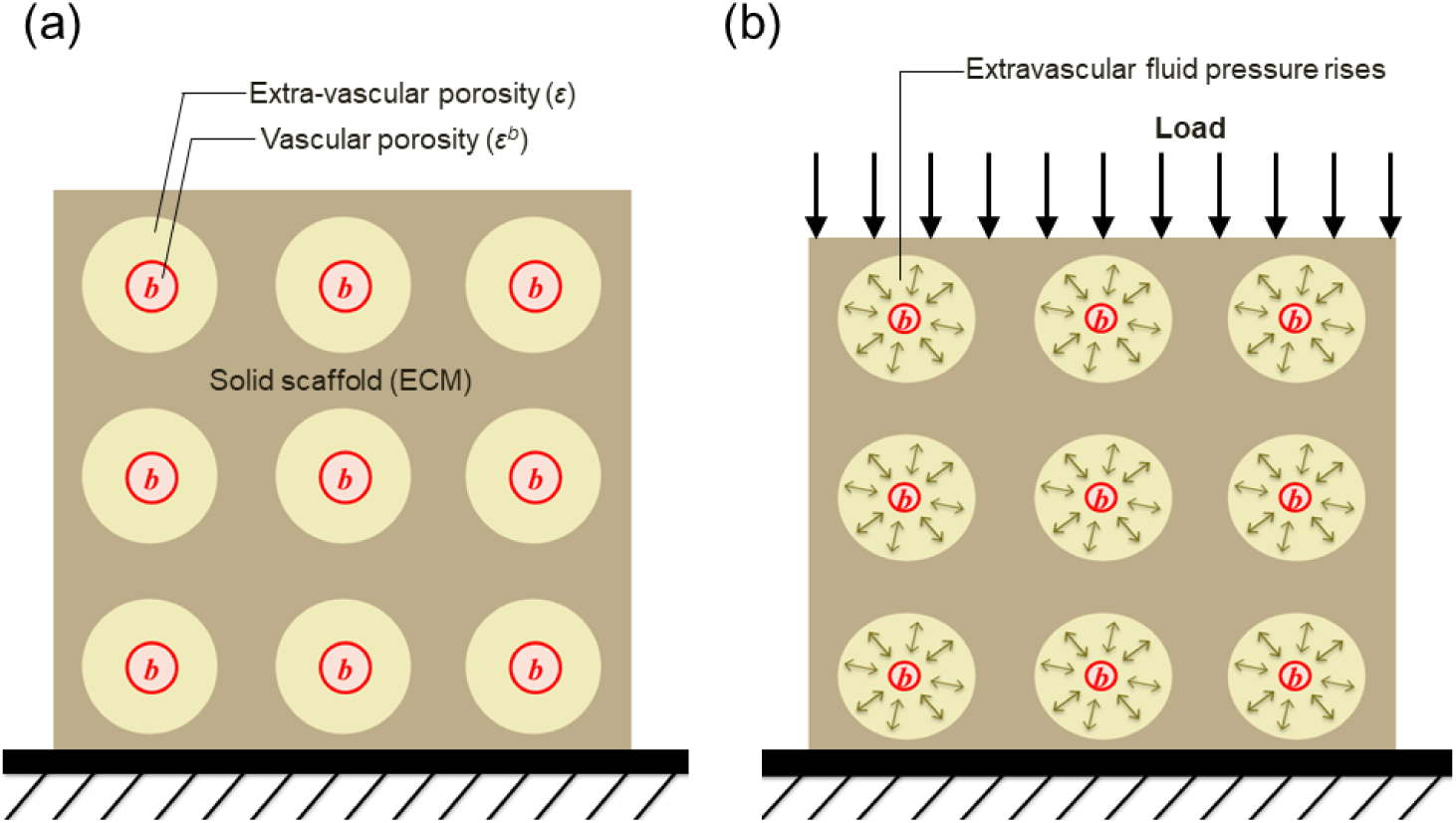
Schematic representation of load transfer. (a) Not loaded configuration. (b) Loaded configuration: load applied to the solid is transmitted to the blood phase thanks to the intercession of the interstitial fluid phase.

### 3.2 Angiogenesis and vascular tumor growth

In this second example tumor growth during the avascular stage, angiogenesis and the subsequent vascular stages are analyzed. A cylindrical domain of 2 *mm* and 0.8 *mm* height is modeled (see figure 7). The tumor spheroid has an initial radius of 30 *µm*. It is placed at 350 *µm* from the bottom of the domain. Two arterioles beds are assumed at the bottom and top basis of the cylinder respectively. Blood pressure in the in the bottom arterioles is 10 Pa higher than that in the top ones so a blood flow is established in the vertical direction from the bottom to the top basis of the cylinder. Near to arterioles a zone of 20 *µm* of thickness is assumed being high vascularized (*ε*^*b*^ = 0.06).

**Fig. 7.**
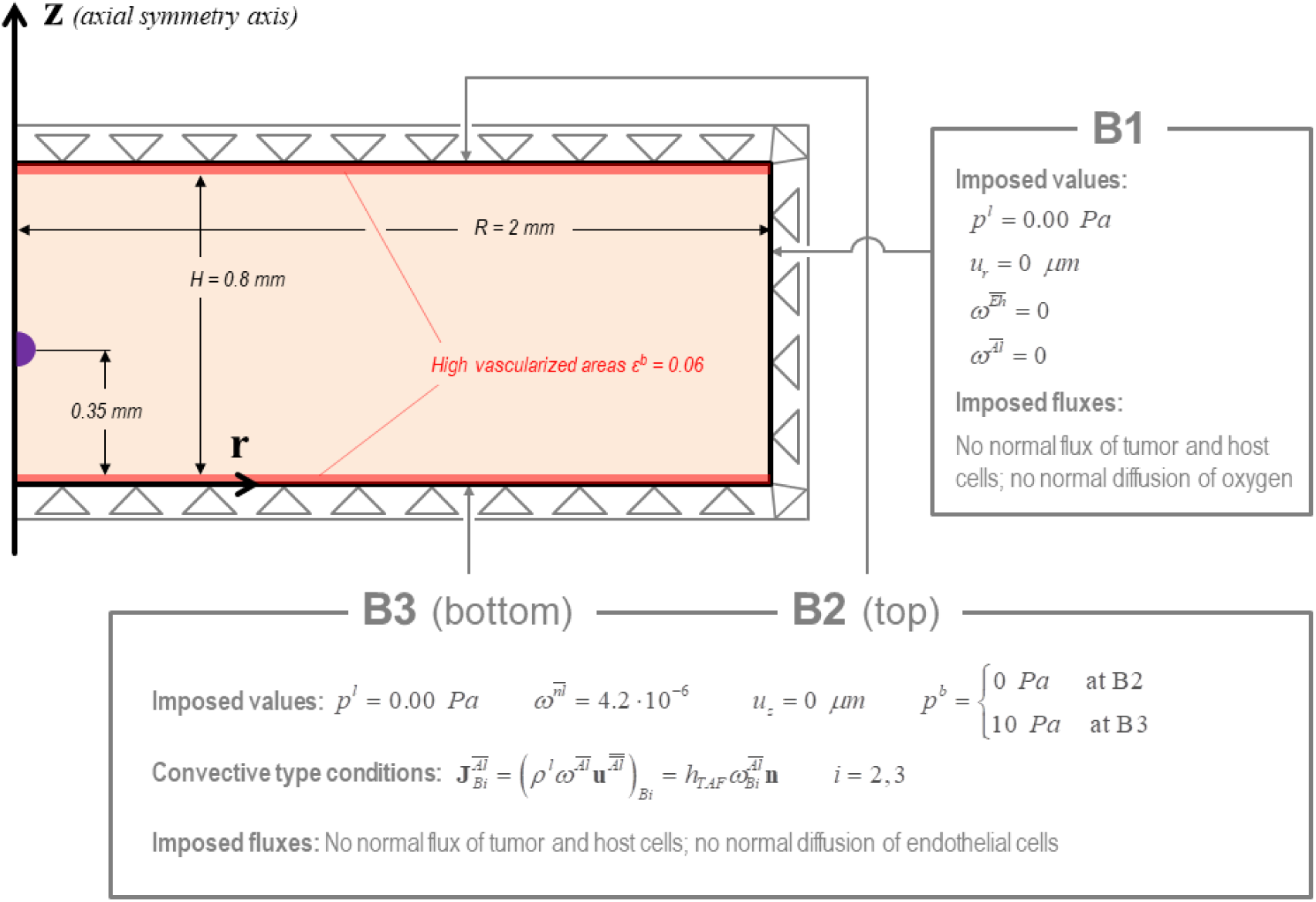
Geometrical configuration and boundary conditions for the second example.

#### 3.2.1 FE mesh, initial and boundary conditions

As for the first example a mixed finite element space is adopted with quadratic elements (triangular with six nodes and quadrangular with eight nodes) used for **u**^*s*^, and linear elements (triangular with three nodes and quadrangular with four nodes) used for scalar primary variables nominally *p*^*th*^, *p*^*hl*^, *p*^*l*^, *p*^*b*^, 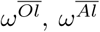 and 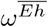. Mass fraction of necrotic tumor cells, 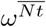, the angiogenesis parameter, *Γ*, and porosity, *ε*, can be treated as internal variables and updated according to variation of primary variables. The computational model exploits cylindrical symmetry.

The system comprises five phases: i) the tumor cells, *t*; ii) the host cells (healthy cells), *h*; iii) a solid ECM, *s*; iv) the interstitial fluid, *l*; and v) blood, *b*.

In the whole discretized domain the extra-vascular porosity is set equal to 0.8, the initial interstitial fluid pressure (IFP), *p*^*l*^, and blood (over)-pressure, *p*^*b*^, are set equal to 0 Pa, while the HC-IF pressure difference, *p*^*hl*^, is set equal to 1800 Pa (this corresponds to a saturation degree of IF, equal to 0.2). Mass fraction of oxygen is initially set equal to 4.2 · 10^*−*6^ (corresponding to dissolved oxygen in the plasma of a healthy individual) while mass fractions of TAF and endothelial cell are null. Volume fraction of blood vessel is set equal 0.001 in the whole computational domain with exception of high vascularized areas (20 *µm* of thickness in proximity of the top and bottom vascular beds) where it is set equal to 0.06. In the tumor zone (purple region in figure 7) all five phases coexist while in the remaining regions of the domain no tumor cells are present. in this zone the initial saturation degree of the tumor cells, *S*^*t*^, is set equal to 0.13, corresponding to *p*^*th*^ ≅ 20 Pa. The initial conditions are summarized in table 3.

**Table 3.** Summary of initial conditions in the three zones of the domain.

| Zone | $p^l$<br>[Pa] | $p^{hl}$<br>[Pa] | $p^{th}$<br>[Pa] | $S^l$<br>[-] | $S^t$<br>[-] | $p^b$<br>[Pa] | $\varepsilon^b$<br>[-] | $\omega^{Ol}$<br>[-] | $\omega^{Al}$<br>[-] | $\omega^{El}$<br>[-] |
| --- | --- | --- | --- | --- | --- | --- | --- | --- | --- | --- |
| Initial tumor zone | 0.0 | 1800 | 20 | 0.2 | 0.13 | 0.0 | 0.001 | $4.2 \cdot 10^{-6}$ | 0.0 | 0.0 |
| Healthy zone | 0.0 | 1800 | 0.0 | 0.2 | 0.0 | 0.0 | 0.001 | $4.2 \cdot 10^{-6}$ | 0.0 | 0.0 |
| Highly vascularized zone | 0.0 | 1800 | 0.0 | 0.2 | 0.0 | 0.0 | 0.06 | $4.2 \cdot 10^{-6}$ | 0.0 | 0.0 |

At the right boundary (B1), Dirichlet essential conditions are assumed for the interstitial fluid pressure, radial displacement, mass fraction of endothelial cells and TAF. For remaining primary variables natural type conditions are assumed (no fluxes of tumor and healthy cells and no diffusion oxygen).

At the top and bottom bounds, boundary conditions are almost identical with exception of the value imposed for blood pressure. Dirichlet essential conditions are assumed for the interstitial fluid pressure, vertical displacement, oxygen mass fraction and blood pressure (assumed at the bottom bound, B3, 10 Pa higher than that of the top bound, B2); a Robin type condition is set for TAF diffusive flow assumed being proportional to boundary concentration of TAF (a coefficient 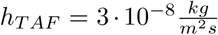 has been identified numerically to obtain a realistic behavior). This condition allows us to model TAF decay due to the presence of the two vascular beds. For remaining primary variables natural type conditions are assumed (no fluxes of tumor and healthy cells and no diffusion of endothelial cells).

The boundary conditions at the *z* axis are assumed respecting cylindrical symmetry. Described boundary conditions are depicted in figure 7.

#### 3.2.2 Derivation of the final form of governing equations

The mathematical model for this second application corresponds to the full version of the multiphase system described in section 2. Dividing each one of equations (17)-(20) by its relative phase density (assumed to be constant), applying the product rule, introducing eqs (27) to express relative velocity of each phase, introducing the previously derived eqn (66) (in eqs (17)-(19) only) and finally expressing *p*^*t*^ and *p*^*h*^ as function of *p*^*l*^ and capillary pressures give

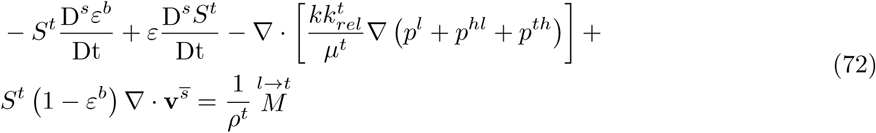

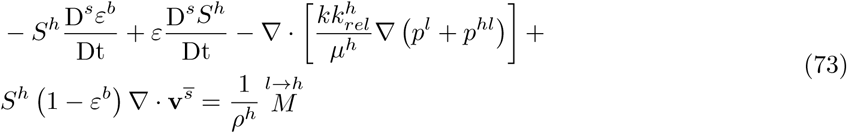

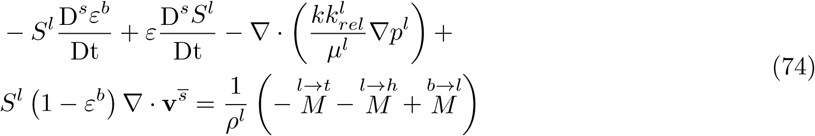

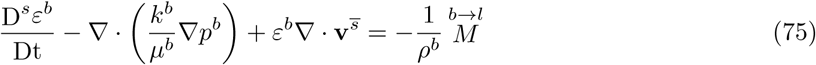

In eqs (72-74) hydraulic conductivity is function of the so-called relative permeability (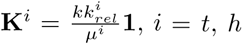 and *l*) to account for the presence of the three fluid phases in the extravascular porosity (14); vascular permeability, *k*^*b*^, is assumed linearly proportional to volume fraction of of blood vessels *ε*^*b*^.

Deriving constitutive eqn (35) with respect to time gives

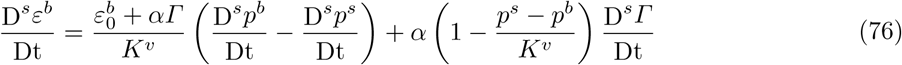

Deriving the solid pressure, eqn (51), with respect to time gives

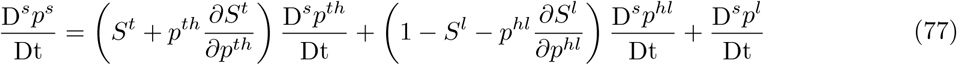

Introducing this equation in eqn (76), this last can be rewritten as

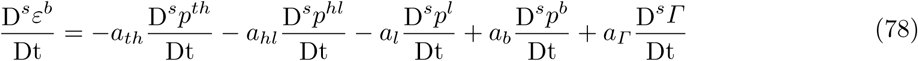

with terms *a*_*i*_ being equal to

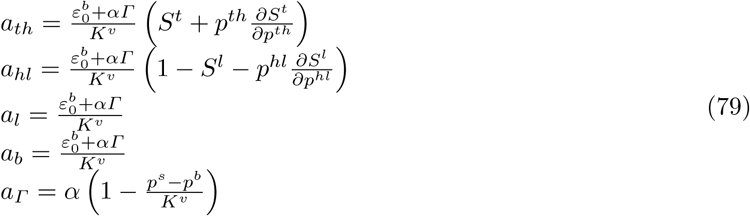

At this point eqn (78) can be introduced in eqs (72-75) which will result expressed in term of the primary variables and internal variable *Γ*. Joining the resulting equations with the rate form of momentum conservation equation of the multiphase system (eqn (26)) and mass balance equations of species, eqs (23-25) (integrated by Fick’s laws defined in subparagraph 2.5.6 and divided by relative phase density), give the final system of governing equation. As for the previous example this system of equation is discretized in time by a centred finite difference scheme. A fix-point scheme is used to address system nonlinearity, with eqn (21) solved at the end of each iteration to updated mass fraction of necrotic tumor cells.

#### 3.2.3 Mass exchange terms related to growth, oxygen consumption and necrosis

Regularized Heaviside functions depending on two constant parameters 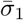 and 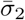 are used for the definition of mass exchange terms. The function 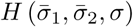 has the following form

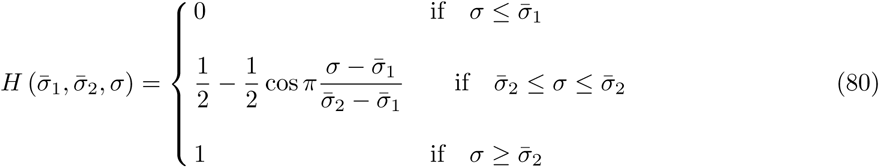

As shown in (12) growth and necrosis and oxygen consumption are importantly impacted by oxygen availability, 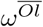, and pressure in the tumor cell phase *p*^*t*^. Two couple of values must be identified for oxygen availability, 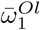 and 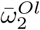, and pressure in the tumor cell phase, 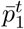 and 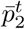, respectively. 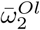 is the value below which cell growth starts to be inhibited by oxygen availability while 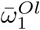 is a critical value below which cell growth is fully inhibited and necrosis starts; conversely 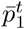 is the critical value of the tumor cell pressure above which growth rate starts to decrease and 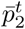 is the critical value above which growth is fully inhibited. Three Heaviside functions based on the four introduced parameters govern growth, oxygen consumption and necrosis

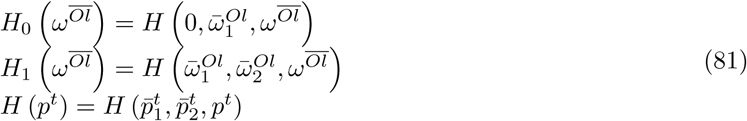

The functions are depicted in figure 8.a-c.

**Fig. 8.**
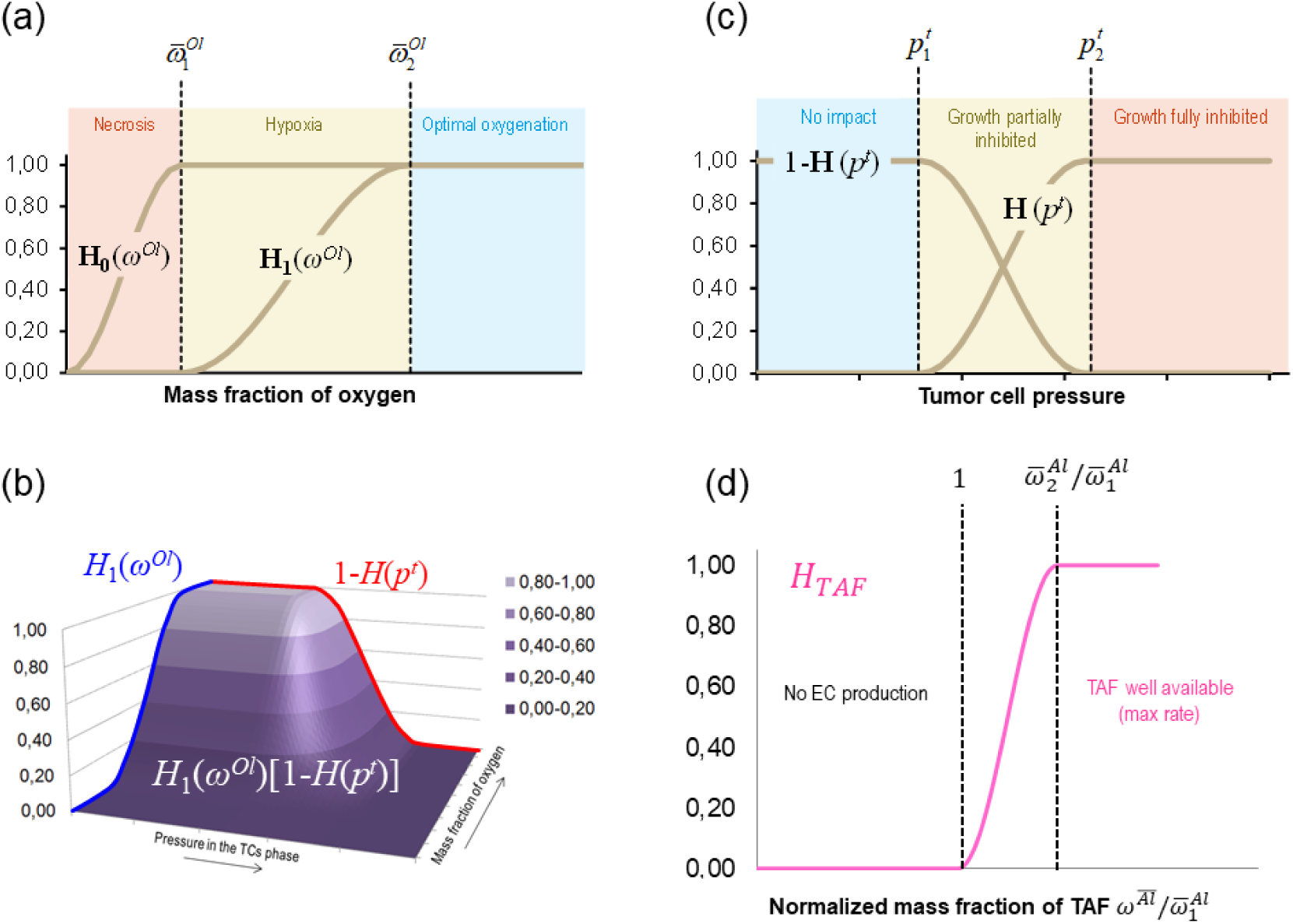
Regularized Heaviside functions governing growth, oxygen consumption, necrosis and endothelial cell production. (a) Impact of oxygen availability on tumor growth; (b) impact of pressure on tumor growth; (c) Coupled effect of tumor pressure and oxygen availability on tumor growth; (d) Function *H*_*TAF*_ versus normalized mass fraction of TAF.

The mass exchange term 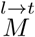 depends on a constant coefficient 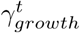, on the functions 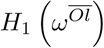 and *H* (*p*^*t*^), and on volume fraction of necrotic tumor cells as follows

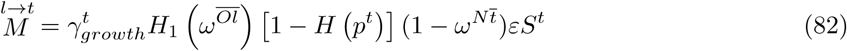

To view coupled impact of functions factors 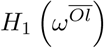 and [1 − *H* (*p*^*t*^)] their product if plotted in figure 8.b.

The sink term accounting for oxygen consumed by tumor cells has the following form

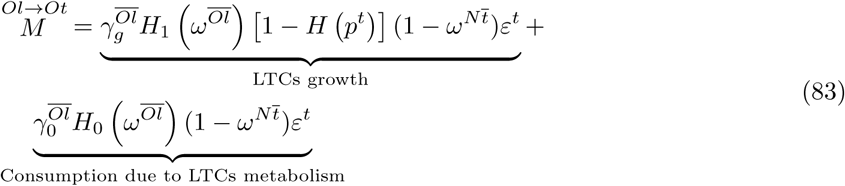

As shown in the previous equation nutrient consumption is due to growth of the tumor cells and to their metabolism. Indeed parameter 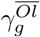 is related to cell division whereas the parameter 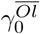 relates to cell metabolism. Oxygen consumed due to cells metabolism depends on the oxygen availability and becomes zero when the mass fraction of oxygen is zero. The function 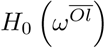 (depicted in figure 8.a) allows us preserving positive values of the local mass fraction of oxygen since negative values have not physical meaning. Observing the form of eqn (83) it easy to shown that, by introducing eqn (82) it can also be expressed in the following form

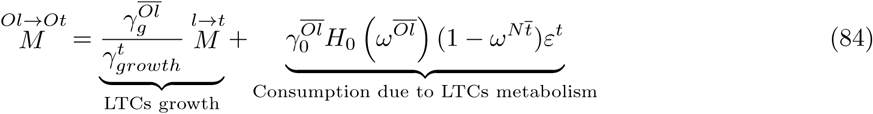

Necrosis depends on nutrient availability and cells pressure. The rate of necrosis, *ε*^*t*^*r*^*Nt*^ in eqn (21), is modeled by the relation

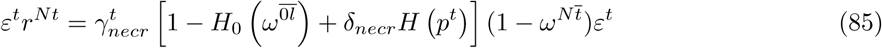

where 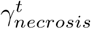 is the rate of cell death in absence of nutrient (and without pressure excess), and *δ*_*necr*_ controls the additional necrosis induced when 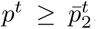. The form of function *H* (*p*^*t*^) (see figure 8.b) indicates that stress impacts cell necrosis rate only when 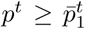. Hence, cell death is assumed to be solely regulated by insufficient concentration of oxygen and excessive mechanical pressure. No drugs or other pro-apoptotic molecules are used in the present model, but eqn (85) can be readily modified to include also this contribution.

The previous presented equations (82), (83) and (85) are an evolution of those used in the the previous version of (12; 14; 15). The advantages of these implemented modifications are: i) the meaning of the coefficients that can easier be related to experiments (for instance 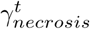 is the rate of cell death in absence of nutrient); ii) the absence of Heaviside step functions which are not suitable from the numerical point of view because often lead to convergence issues.

#### 3.2.4 Mass exchange terms related to TAF release and angiogenesis

TAF is released by hypoxic cells. The higher is hypoxia, the higher is TAF release rate. The following equation is proposed here

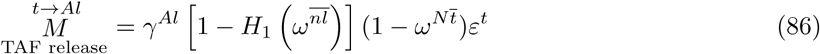

where *γ*^*Al*^ is a constant parameter representing maximum release rate. Production of endothelial cells is related to the intensity of the TAF signal. The higher is the mass fraction of TAF the higher is production rate of endothelial cells. To model this dependence a new regularized Heaviside function is introduced

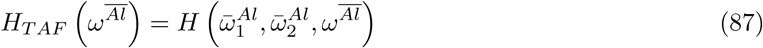

This function is depicted in figure 8.d. If mass fraction of TAF is lower than the critical value 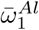 endothelial cells are produced, when mass fraction of TAF is comprised between 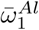 and 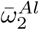 endothelial cell generation depend on the intensity of the TAF signal, finally when TAF signal is relatively high (mass fraction higher than 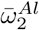) endothelial cells are produced at the maximum rate. The rate of production of endothelial cells depends also on the degree of vascularization of the tissue. The proposed equation reads

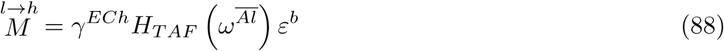

Finally the exchange of oxygen from the vascular porosity, *ε*^*b*^, to the extravascular porosity, *ε*, is modeled with a convective type condition at the interface between two porosities. A coefficient *h*_*v*_ is representative of vessel wall permeability. The following relationship is adopted

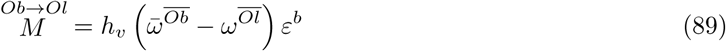

where 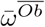 is the mass fraction of oxygen in the blood assumed as a constant parameter.

#### 3.2.5 Results

Parameters used for the numerical simulation are reported in the following four tables: table 4 shows parameters associated to cell phases and IF; table 5 refers to those related to oxygen, TAF and EC diffusion; table 6 shows parameters associated to the ECM; finally table 7 regroups parameters associated to the vascular porosity, blood and angiogenesis.

**Table 4.** Parameters depending on cells’ type and IF.

| Parameter | Symbol | Value | Unit |
| --- | --- | --- | --- |
| Dynamic viscosity of IF | $\mu^l$ | 0.01 | Pa.s |
| Dynamic viscosity of TC | $\mu^t$ | 36 | Pa.s |
| Dynamic viscosity of HC | $\mu^h$ | 36 | Pa.s |
| Critical mass fraction of oxygen | $\bar{\omega}_1^{Ol}$ | $1 \cdot 10^{-6}$ | - |
| Inhibition mass fraction of oxygen | $\bar{\omega}_2^{Ol}$ | $3 \cdot 10^{-6}$ | - |
| Growth coefficient of tumor cells | $\gamma_{growth}^t$ | $9.6 \cdot 10^{-3}$ | kg/(m<sup>3</sup>.s) |
| Necrosis coefficient | $\gamma_{necrosis}^t$ | $9.6 \cdot 10^{-3}$ | kg/(m<sup>3</sup>.s) |
| Consumption related to growth | $\gamma_{growth}^{Ol}$ | $2.4 \cdot 10^{-4}$ | kg/(m<sup>3</sup>.s) |
| Consumption related to metabolism | $\gamma_0^{Ol}$ | $6 \cdot 10^{-4}$ | kg/(m<sup>3</sup>.s) |
| HC-IF interfacial tension | $\sigma_{hl}$ | 72 | mN/m |
| TC-HC interfacial tension | $\sigma_{th}$ | 12 | mN/m |

**Table 5.** Parameters related to oxygen, TAF and endothelial cells diffusion.

| Parameter | Symbol | Value | Unit |
| --- | --- | --- | --- |
| Diffusion coeff. of oxygen in IF | $D_0^{Ol}$ | $3.2 \cdot 10^{-9}$ | m<sup>2</sup>/s |
| Diffusion coeff. of TAF in IF | $D_0^{Al}$ | $2.9 \cdot 10^{-11}$ | m<sup>2</sup>/s |
| Coefficient $\delta_i$ ($i = O, A$) in eqn (54) | $\delta_i$ | 2 | - |
| Diffusion coeff. of EC in HC | $D_0^{Eh}$ | $1 \cdot 10^{-13}$ | m<sup>2</sup>/s |
| Chemotactic coefficient in eqn (55) | $C$ | $1 \cdot 10^{-4}$ | - |
| Normal mass fraction of oxygen | $\bar{\omega}_{env}^{Ol}$ | $4.2 \cdot 10^{-6}$ | - |

**Table 6.** Parameters depending on ECM type.

| Parameter | Symbol | Value | Unit |
| --- | --- | --- | --- |
| Poisson's ration of ECM | $\nu$ | 0.4 | - |
| Young's modulus of ECM | $E$ | 1000 | Pa |
| Extravascular porosity (initial) | $\varepsilon$ | 0.8 | - |
| Coefficient $a$ in eqs (49,50) | $a$ | 590 | Pa |
| Intrinsic permeability | $k$ | $1.8 \cdot 10^{-15}$ | m<sup>2</sup> |

**Table 7.** Vasculature, angiogenesis and blood parameters.

| Parameter | Symbol | Value | Unit |
| --- | --- | --- | --- |
| Vascular compressibility parameter | $K^v$ | 5000 | Pa |
| Microvascular intrinsic permeability | $k^b$ | $5 \cdot 10^{-16}$ | $\text{m}^2$ |
| Blood dynamic viscosity | $\mu^b$ | 0.004 | Pa·s |
| TAF release coefficient | $\gamma^{Al}$ | $1 \cdot 10^{-10}$ | $\text{kg}/(\text{m}^3 \cdot \text{s})$ |
| Mass fraction of TAF activating EC proliferation | $\bar{\omega}_1^{Al}$ | $8 \cdot 10^{-12}$ | - |
| Mass fraction of TAF maximizing EC proliferation | $\bar{\omega}_2^{Al}$ | $8.8 \cdot 10^{-12}$ | - |
| Production rate of EC | $\gamma^{Eh}$ | $1 \cdot 10^{-4}$ | $\text{kg}/(\text{m}^3 \cdot \text{s})$ |
| Parameter governing the rate of angiogenesis | $b_\Gamma$ | 50 | - |
| Additional vascular porosity when $\Gamma = 1$ | $\alpha$ | 0.06 | - |

To simplify model parametrization and results interpretation the effect of pressure on tumor cell metabolism-growth-necrosis and oxygen mass exchange terms from the vascular to the extra-vascular compartment (terms 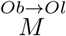 and 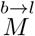) are not accounted (so associated parameters are not reported in the tables). Figures 9 and 10 show volume fraction of blood vessels (left column) and mass fraction of oxygen (right column) before and after angiogenesis respectively. Furthermore the two white lines in the left columns of figures 9 and 10 depict the tumor front and the limit between fully viable areas and more internal zones where necrosis occurs while the three isovalue lines in the right columns refers to TAF mass fraction. In particular the yellow isovalue line is the critical mass fraction of 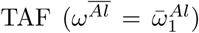 that activates endothelial cells production.

**Fig. 9.**
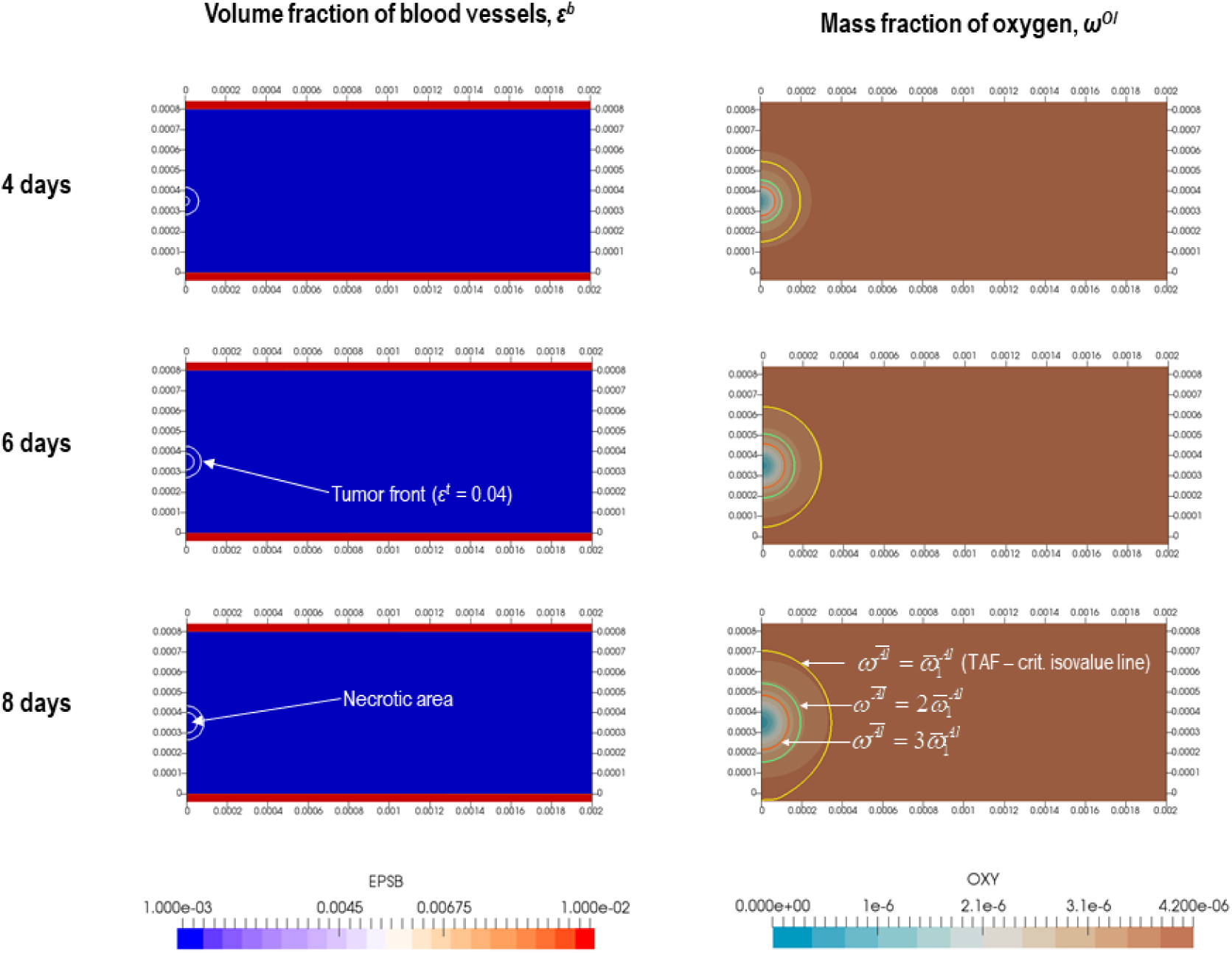
Volume fraction of blood (left column) and mass fraction of oxygen (right column) prior to angiogenesis (from 0 to 8 days).

**Fig. 10.**
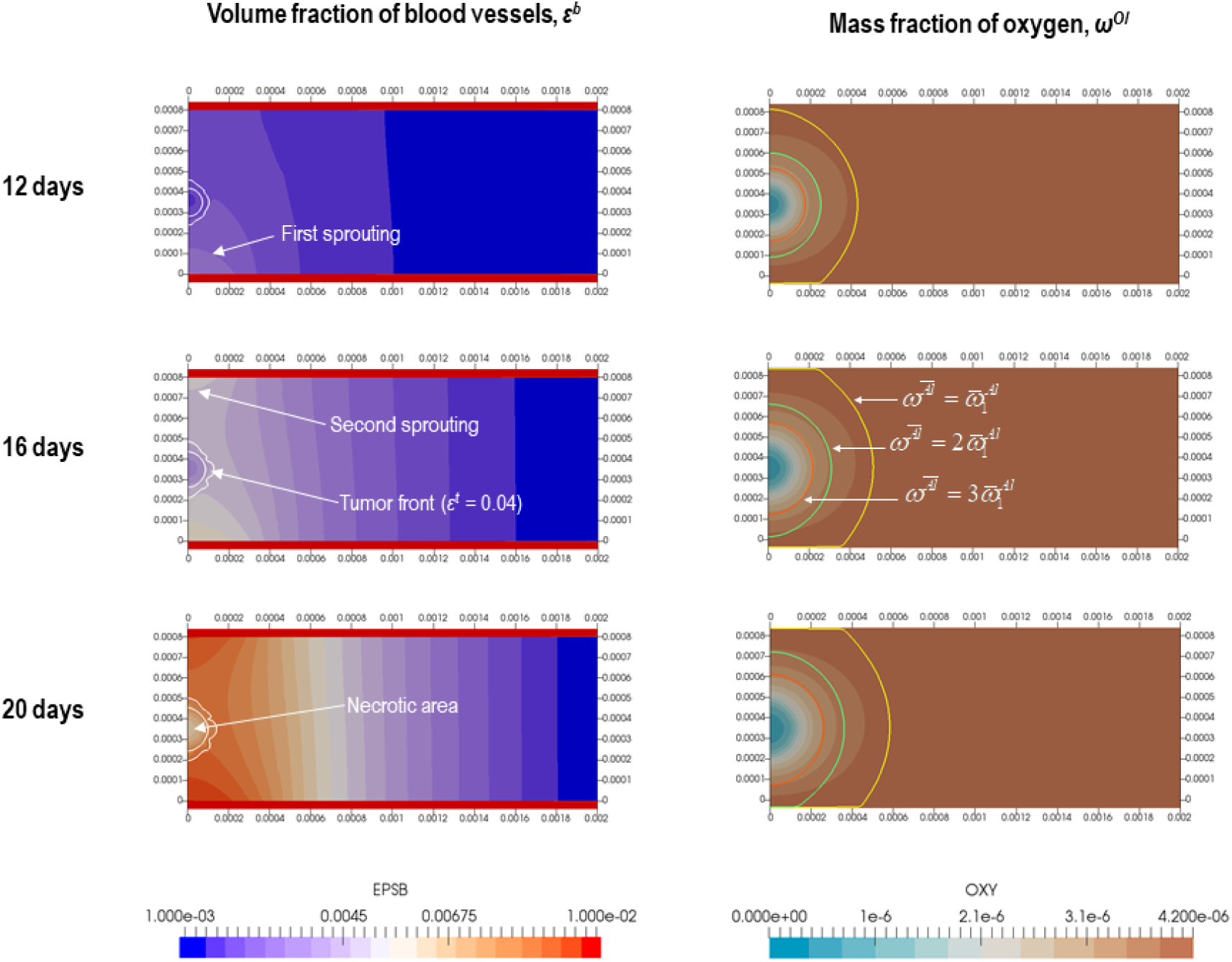
Volume fraction of blood (left column) and mass fraction of oxygen (right column) during angiogenesis process (from 12 to 20 days).

Figure 9 shows that during the first phase tumor growth in a quite spherical shape and an hypoxic area progressively appears in the center of the tumor spheroid. Such hypoxic area starts to necrotize and to release TAF. TAF mass fraction increases with time (see the three isovalue lines in the right part of figure 9). The tumor is more in proximity of the bottom vascular bed so the TAF critical mass fraction is reached firstly at the bottom vascular bed (see yellow line at 8 days in figure 9). This induces proliferation of endothelial cells that start to migrate and to form the new microvasculature. A first sprout is clearly visible in figure 10 at 12 days (left column). At this time TAF mass fraction reaches it critical value at the top vascular bed also (yellow isovalue line in the right column), and four days later a second sprout becomes visible (see figure 10, left column, 16 days). As depicted in figure 10, at 20 days the new vessels surround the tumor mass that has assumed a more complex growth pattern due to the modifications and spatial heterogeneities induced by neo-vascularization in its surrounding.

## 4 Conclusions

In this paper the tumor growth model of (15) is further extended to account for angiogenesis and growth in the vascular stage. Improvements are substantial and allow us to define a hierarchy of mathematical models (see figure 2) of decreasing complexity: the most general vascular growth model with angiogenesis (VGA); a simplified vascular growth model (VG) (without angiogenesis); and finally the avascular growth model (AG) (corresponding to the model of (15)). The paper shows how the biophysical behavior of the multiphase vascularized tissue (tumor growth, blood flow, angiogenesis, etc.) can be modeled in a mechanistic fashion within the framework of porous media mechanics thanks to a non-conventional *ad hoc* definition of the effective stress tensor. The mathematical model allows us to account for the bio-chemo-mechanical interaction between the five phases constituting the tissue: the ECM, the two cell populations, the interstitial fluid and blood. Blood is in the vasculature while cell phases and interstitial fluid are in a separate porous compartment. ECM is the solid fibrous/porous scaffold. The mathematical model is detailed in the first part of the paper then two applications are presented.

The first application, “one-dimensional bio-consolidation: one fluid + blood”, is very simple but at the same time tremendously useful to understand how the model works. It is shown that when an external load is applied it is instantaneously transferred to the fluid in the extravascular porosity, which transfers it to the blood phase: both extravascular fluid and blood start to move and to be drained but with a different dynamics (see figures 4 and 5). To better explain this behavior, one can observe from the blood volume fraction constitutive model (eqn (35)) that the solid scaffold and blood in the vasculature are mechanically connected thanks to the intercession of the extravascular fluid (see idealized mechanism in figure 6). Considering an indirect coupling between the solid scaffold (ECM) and the blood allows us to substantially simplify the closure of the mathematical model otherwise difficult to perform without the introduction of unreliable simplifications declassing the biophysical significance of the model (*e.g.* incompressible vessels, constant blood pressure, etc.). Biophysical significance is conversely not prejudiced by the assumption of an indirect ECM-blood interaction. Actually, in biological tissues the volume fraction of the ECM and that of blood vessels are normally much lower than the cumulative volume fraction of TC, HC and IF; hence, both ECM and blood vessels have privileged mechanical relationships with extravascular fluid phases (interaction modeled by means of the solid pressure *p*^*s*^) and not between themselves.

In the second application the results of the full model are presented. A tumor mass growing between two vascular beds is modeled. At a certain time the oxygen mass fraction in the core of the tumor decreases below a critical value inducing necrosis and production of tumor angiogenic factors (TAF). TAF diffuse within the tissue, so TAF mass fraction increases progressively until it starts to stimulate division of endothelial cells in the highly vascularized vascular beds. Endothelial cells proliferate and move towards the tumor driven by their own gradient and by the gradient of TAF (*chemotaxix*). Endothelial cells move and at the same time arrange themselves forming the microvasculature and improving blood flow in the neo-vascularized areas. As anticipated in the introduction, some of adopted angiogenesis-constitutive relations are not of demonstrated biophysical significance because no specific information is available in literature for their validation. Hence, results of the full VGA model have today only a qualitative value because are not validated with a suitable *in vitro* or *in vivo* experiment. Taking advantage from a previous *in vitro*-*in silico* study (20) where we have exhaustively validated the AG model, an improvement of the presented experimental methodology (confined co-culture of endothelial and tumor cells based on Cellular Capsule Technology, (21)) is under development to validate VG and VGA models.

Despite the model is here specialized and applied for numerical simulation of vascular tumor growth, this general framework can be also used to simulate drugs delivery, physiological behavior of healthy tissues and further extended to account for the lymphatic system.

## A Appendix: Chemotactic-fickian model for EC diffusive velocity

Let us assume that all non-endothelial cell species (*e.g.* non endothelial host cells, chemical species etc.) can be modeled jointly as a species named *H*, dominant species in *h*. Consequently let us treat species *E* as a diluted species in *h*. For such a situation, neglecting body forces potential and assuming isothermal condition, TCAT (11) provides from simplified entropy inequality the following force-flux pair

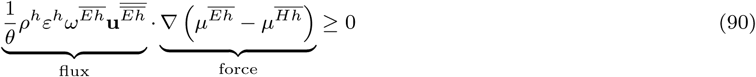

where *θ* is the temperature and 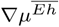 and 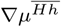 are chemical potentials of endothelial and non-endothelial species respectively. At equilibrium both force and flux terms are zero. Conversely near equilibrium a first order closure approximation for the force-flux relationship reads

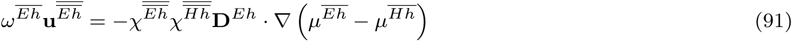

where **D**^*Eh*^ is a second order symmetric tensor, and 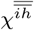 is the molar fraction of species *i*. Gibbs-Duhem equation for this binary mixture reads

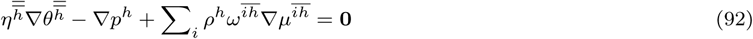

The expected pressure gradient in the phase *h* is relatively weak. Hence, for the isothermal case considered eqn (89) reduces for a binary mixture to

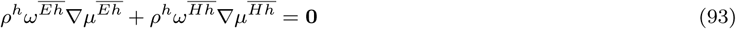

Eqn (90) allows us to obtain the gradient of the chemical potential of species *H* as

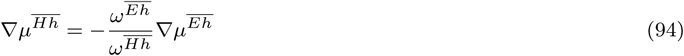

being 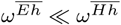 it follows that 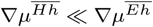 and consequently eqn (90) reduces to

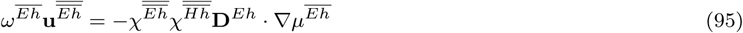

To gain usefulness of the previous equation a relationship between the macroscale chemical potential of species *E* and its mass fraction is needed. The macroscale chemical potential for the species *E* can be written as

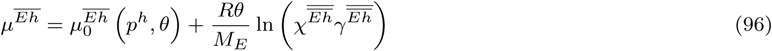

where 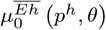 is a reference chemical potential for species *E, R* is the ideal gas constant, *M*_*E*_ is the molar mass of species *E*, and 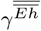 is the macroscale activity coefficient. Being the system in isothermal condition (*θ* = *θ*_0_) and the impact of pressure gradient of phase *h* assumed negligible, differentiating this expression in space gives

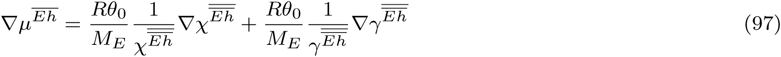

For dilute species the macroscale activity coefficient is usually assumed constant and equal to 1. To account for chemotaxis we set here an activity coefficient linearly dependent on the mass fraction of TAF in *h*. We assume here local chemical equilibrium so despite the mass fraction of TAF in *h*, 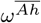, is not a primary variable of the model, its value can be linearly related (as a first approximation) to mass fraction of TAF in the adjacent IF phase, 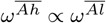. This allows us to assume that the activity coefficient, 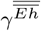, is linearly dependent on the mass fraction of TAF in *l*. Thanks to short-range diffusion and molecular signalling the TAF in the phase *l* interferes (*via* the *hl* interface) with endothelial cell modifying their activity coefficient. The following relationship is assumed with *c* constant chemotactic coefficient and 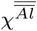 the molar fraction of TAF in IF

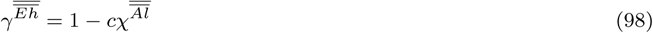

Introducing eqn (98) in eqn (97) gives

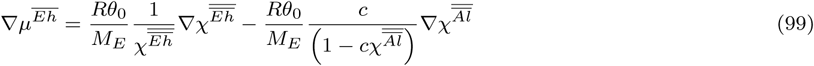

We reasonably assume here that molar masses of phases *h* and *l* are weakly affected by variation of species concentration. This allows us to assume constant molar masses of phases *h* and *l* and to express molar fraction of species *E, A* and *H* as function of respective mass fractions

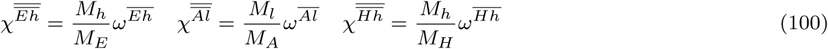

Introducing the first two relationships of eqn (100) in eqn (99) and setting 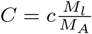 give

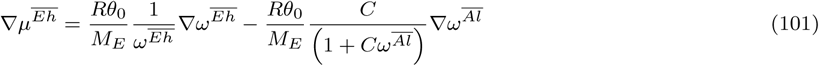

We now introduce eqn (101) in eqn (95) and express 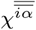 as function of 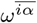. After some calculations we obtain

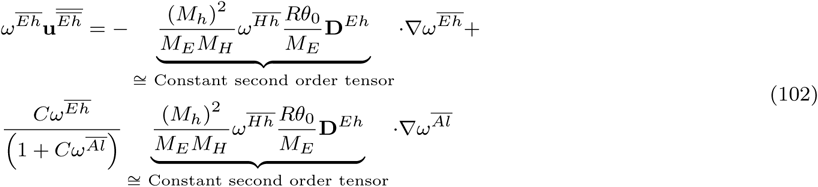

As shown in the previous eqn some quantities are expected to stays almost constant being always 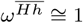. This observation allows us to rewrite the previous equation in a simplified form

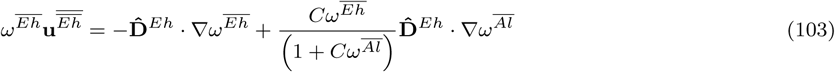

Where the diffusivity tensor 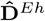 reads

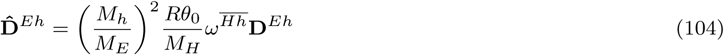

We assume here an isotropic effective diffusivity which linearly increases with volume fraction of phase *h*. Therefore eqn (103) can be rewritten in this form

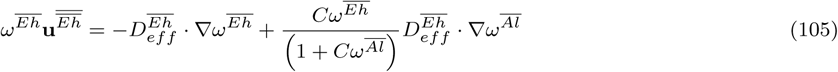

where indicating with 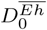 the bulk diffusivity of endothelial cells in *h*, the effective diffusivity reads

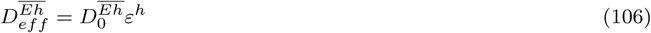

